# Selection on the fly: short term adaptation to an altered sexual selection regime in *Drosophila pseudoobscura*

**DOI:** 10.1101/2022.03.16.484639

**Authors:** Carolina Barata, Rhonda R. Snook, Michael G. Ritchie, Carolin Kosiol

**Author notes:** Joint corresponding authors and.

## Abstract

Experimental evolution studies are powerful approaches to unveil the evolutionary history of lab populations. Such studies have shed light on how selection changes phenotypes and genotypes. Most of these studies have not examined the time course of adaptation under sexual selection manipulation, by resequencing the populations’ genomes at multiple time points. Here, we analyse allele frequency trajectories in *Drosophila pseudoobscura* where we altered their sexual selection regime for 200 generations and sequenced pooled populations at 5 time points. The intensity of sexual selection was either relaxed in monogamous populations (M) or elevated in polyandrous lines (E). We present a comprehensive study of how selection alters population genetics parameters at the chromosome and gene level. We investigate differences in the effective population size – *N*_*e*_ – between the treatments, and perform a genome-wide scan to identify signatures of selection from the time-series data.

We found genomic signatures of adaptation to both regimes in *D. pseudoobscura*. There are more significant variants on E lines as expected from stronger sexual selection. However, we found that the response on the X chromosome was substantial in both treatments, only more marked in E and restricted to chromosome arm XR in M. *N*_*e*_ is lower on the X at the start of the experiment, which might indicate a swift adaptive response at the onset of selection. Additionally, we show that the third chromosome was also affected by elevated polyandry. Its distal end harbours a region showing a strong signal of adaptive divergence in E lines.

## 1 Introduction

Evolutionary biologists have put considerable effort into uncovering how social environments shape evolution, especially those that change sexual selection pressures. Studies over the years have found differences in courtship phenotypes as well as other fitness-related traits caused by altered mating systems (Chapman et al., 1995; Wigby and Chapman, 2004; Chenoweth and Blows, 2005; Hollis et al., 2017). Due to the effects of mate competition, male harm has also been found to evolve under specific environmental conditions (Holland and Rice, 1999; Yun et al., 2019).

A few key studies have tried to identify the genetic basis of adaptation to a new sexual selection regime. Sexually antagonistic loci were initially hypothesised to be more prevalent on the X chromosome (Rice, 1984). This prediction was confirmed in several studies (e.g. Chippindale et al., 2001; Innocenti and Morrow, 2010). However, subsequent theoretical predictions suggested that autosomes are just as likely to harbour sexually antagonistic polymorphism as the X under certain conditions (Fry, 2009; Ruzicka and Connallon, 2020). Others have subsequently shown that sexual selection seems to affect many of the same genomic regions as those affected by natural selection regardless of chromosomal location (Chenoweth et al., 2015). More surprisingly, the X chromosome was found not to be a hotspot for sexually antagonistic variation in lines of *Drosophila melanogaster* (Ruzicka et al., 2019). We also know that the resolution of sexual conflict over gene expression optima is involved in the response to sexual selection (Innocenti and Morrow, 2010). Evidence indicates that sexual antagonism can lead to sex-biased gene expression within a relatively short timescale (Wright et al., 2018). The importance of sexual selection in shaping the genomic landscape of a population is therefore still largely undiscovered. We will characterise the adaptive response of polymorphic sites throughout the genome. It will address some of general patterns in short-term adaptation that have started to emerge. In particular, we will focus on whether the response to an altered mating system is predominantly on the X chromosome or more evenly distributed along the genome.

Here, we investigate patterns of genetic adaptation of *Drosophila pseudoobscura* flies in a socially manipulated environment across 200 generations of evolution. The experiment consisted of rearing replicated populations under either monogamy – M – or elevated polyandry – E. These two treatments should relax or increase sexual selection, respectively. It has been shown that behavioural and physiological traits have diverged between these lines throughout the experiment. These include courtship song and male mating and courtship rates. In summary, E males produced more attractive song, show decreased singing latency and faster songs over longer periods of time (Snook et al., 2005; Debelle et al., 2017). These males also had higher courtship rates (Crudgington et al., 2010). In contrast, M males had smaller accessory glands and sired fewer progeny (Crudgington et al., 2009). Interestingly, female preference also seems to have coevolved with male signal in opposite directions between the two selection regimes (Debelle et al., 2014).

It was thus demonstrated that sexual selection substantially affected multiple traits as populations adapted. However, a better understanding of the genetic mechanisms responsible for differences in phenotype is needed. Analyses of gene expression patterns in virgin M and E females showed that 14% of the transcriptome was differentially expressed (Immonen et al., 2014). In addition, 70% of these differences were sex biased. This suggests that loci under sexually antagonistic selection might be contributing to divergence between the treatments. Nevertheless, more evidence is required to make a compelling case for sexual conflict as a major driver of adaptation in the E line flies. The prediction that sexual selection should increase the number of male-biased genes was examined (Veltsos et al., 2017). Both female and male transcriptomes were sequenced and revealed that the majority of differentially expressed genes was found in males’ heads, which is consistent with the importance of behavioural traits. Conversely, M treatment flies were predicted to exhibit a feminisation of the transcriptome. In M populations, there was indeed a feminisation of male heads but, contrary to expectations (Haig, 2006; Hollis et al., 2014), male abdomens and both female heads and abdomens were masculinised. This is important since the abdomens house the sex-specific reproductive tissues.

Transcriptome evolution therefore seems to be a large part of the adaptive response to sexual selection. However, it is still unclear how sexual selection can cause allele frequencies to change in the short-term. Genomic time-series data can provide a missing link between phenotypic changes and proof of selection acting on the genome. For this reason, investigating allele frequency trajectories alongside experimental phenotypes in an Evolve & Resequence (E&R) design can prove very useful. They can help determine the rate and strength of selection driving genomic responses. E&R studies focus on adaptation from standing genetic variation and can help reveal signatures of selection by investigating allele frequency changes. Experimental populations are typically sampled and resequenced repeatedly within a certain number of generations. Samples at two time points can be used to test for selection by finding allele frequency changes that differ significantly between treatments (e.g. Pearson’s chi-square, *χ*^2^, test as in Griffin et al., 2017; Fisher’s Exact test as in Burke et al., 2010; or the Cochran-Mantel-Haenszel, CMH, test as in Barghi et al., 2017). However, such approaches lack the ability to take advantage of the allele’s frequency trajectory. In contrast, more probabilistic modelling frameworks use time-series data to fully describe frequency trajectories. In particular, time-series approaches gain a lot from accounting for sampling noise typical of E&R experimental designs.

There are theoretical predictions on the genetic basis of adaptation to an altered mating system that we can consider. First, diversity on the autosomes (A) is expected to differ from X-linked diversity due to differences in the effective population size. Under monogamy, X/A diversity ratios are predicted to be roughly 3*/*4. Under polyandry, however, these ratios are expected to shift towards even lower values especially if populations are founded following a bottleneck (Pool and Nielsen, 2008). This effect should perhaps be counter-acted by the experimental design in our study. The family size for E and M populations was set to ensure that *N*_*e*_ on the autosomes was roughly the same in both lines (addressed in Snook et al., 2009). In addition, if most beneficial mutations on the X chromosome are partially recessive, diversity on the X is predicted to be lower compared to that on the autosomes (Betancourt et al., 2004). These effects combined with sexual selection pressures are expected to result in a marked reduction in diversity on the X chromosome. The X chromosome is known to harbour genes important for mating success, namely accessory gland proteins (Acps) which are involved in sperm production along with other seminal fluid proteins.

With the appropriate statistical framework, we can now characterise allele frequency changes caused by sexual selection in these *D. pseudoobscura* populations. Here, we looked for evidence of adaptation in M and E line females both at the chromosome and gene level. Our study builds on the work of Wiberg et al. (2021) who examined genomic variation between the two treatments after ≈ 160 generations of selection. Wiberg et al. (2021) found “islands” of differentiation between the lines located on the X and 3^rd^ chromosomes. This work offers a more comprehensive analysis of full allele frequency trajectories. Whilst resequencing early generations was not possible, we produced and analysed a pool-seq time-series consisting of five time points throughout those 15 years of evolution in the lab. Starting at generation 21, this time-series allows us to better understand both short-term and long-term effects. Our study assumes that adaptation has proceeded from standing genetic variation in these populations so that the effect of new mutations is negligible. We estimated the effective population size – N_e_ – which will be influenced by census size, mating system, and strength of selection for the four main *D. pseudoobscura* chromosomes: 2, 3, 4 and X. We used a Bayesian modelling approach – Bait-ER (Barata et al., 2020) – to infer selection on individual SNPs that allows for finding potential targets of selection. We combined individual SNP tests for selection with window-based estimates of the effective population size, *N*_*e*_, which gave us a clearer view of the process of adaptation. We found that there was a substantial response to selection with the two treatments differing in their rate of adaptation.

## 2 Results

### 2.1 Diversity and allele frequency changes

We first investigated the allele trajectories by looking at allele frequency spectra throughout the experiment. The time-series consists of five time points from generation 21 to generation 200 (T1: 21-22; T2:59-63; T3:112-116; T4:160-164; T5:200; see **table S1** for more details). Frequency spectra at the start are flat distributions with maxima roughly at 0.5-0.6 (**fig. 1**). Alleles fixed at high rates, with the most fixations between time point 3 and time point 4 for M lines and time points 2 and 3 for E lines. Up to 29.5% and 16.9% more fixed sites than in the previous time point were observed for M and E, respectively. This indicates that diversity was more swiftly reduced in E populations, as expected if sexual selection is stronger in this regime. Allele frequency changes between first and last time point show distributions that are highly skewed towards low values (**fig. S7**). This is especially true in the case of the X chromosome.

**Figure 1:**
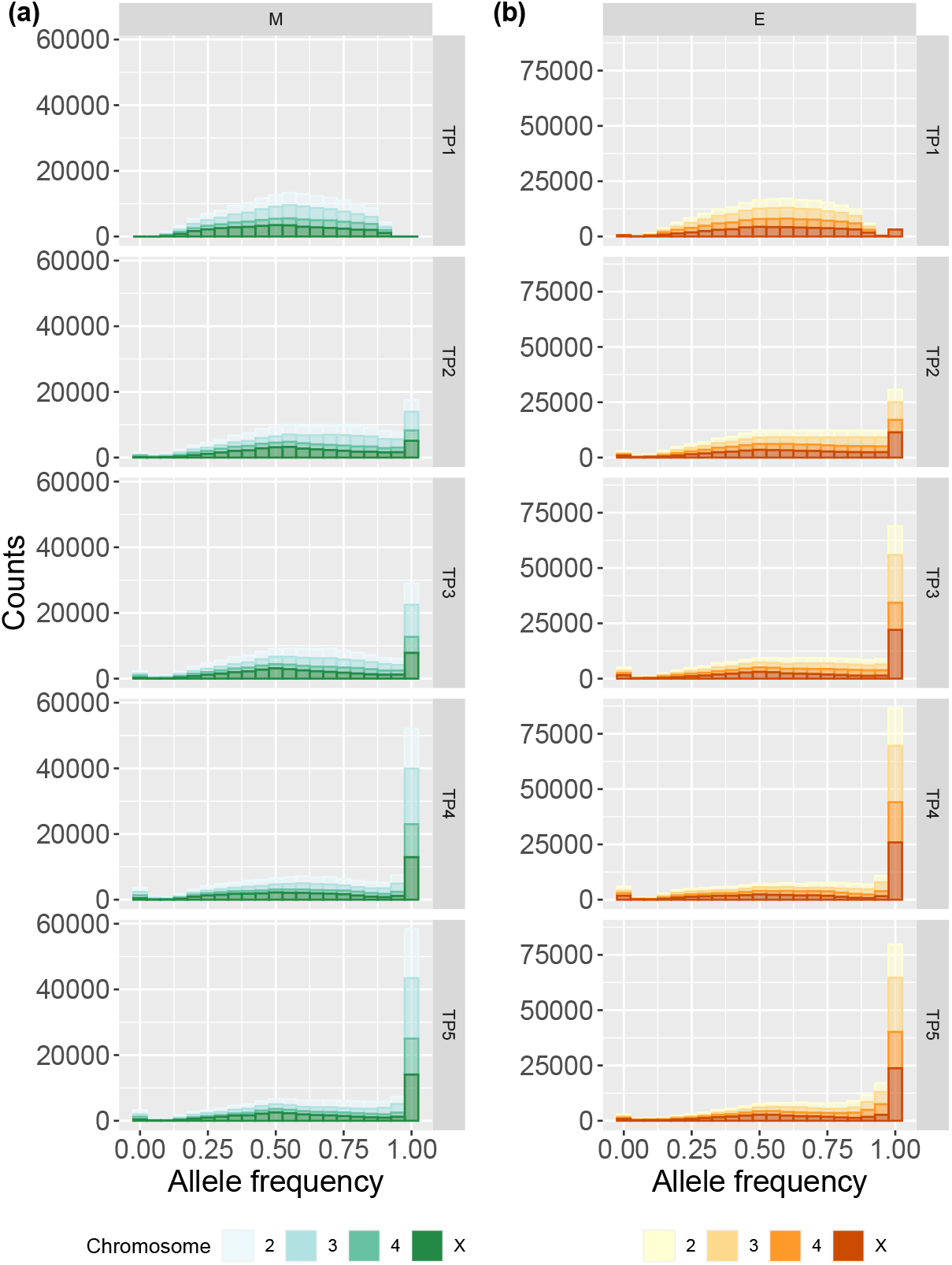
Allele frequency spectra in M and E populations. Allele frequency spectra in (a) for M and (b) for E populations per time point (rows) for each replicate population (columns). Each chromosome is coloured in a different shade of green (M) or orange (E) as seen on the bottom legend.

Nucleotide diversity was measured in 250kbp windows for each chromosome separately and at each time point. Diversity distributions show a marked reduction as time passes, particularly from time point 1 to time point 2 (**fig. 2**). All chromosomes’ densities peak at 0.4-0.5 per site at the first time point. At the end, densities for chromosomes 3 and X flatten out, especially for E flies. Interestingly, in M lines, *π* on the 3^rd^ chromosome becomes skewed towards very low values in later generations. In contrast, chromosomes 2 and 4 maintain more diversity. Taken together, these results suggest that selection was pervasive on the 3^rd^ and X chromosomes resulting in more windows of very low *π* across treatments. Window estimates along each chromosome exhibit some diversity peaks, particularly on the 3^rd^ chromosome, that become flat towards the end of the experiment (**fig. S8**).

**Figure 2:**
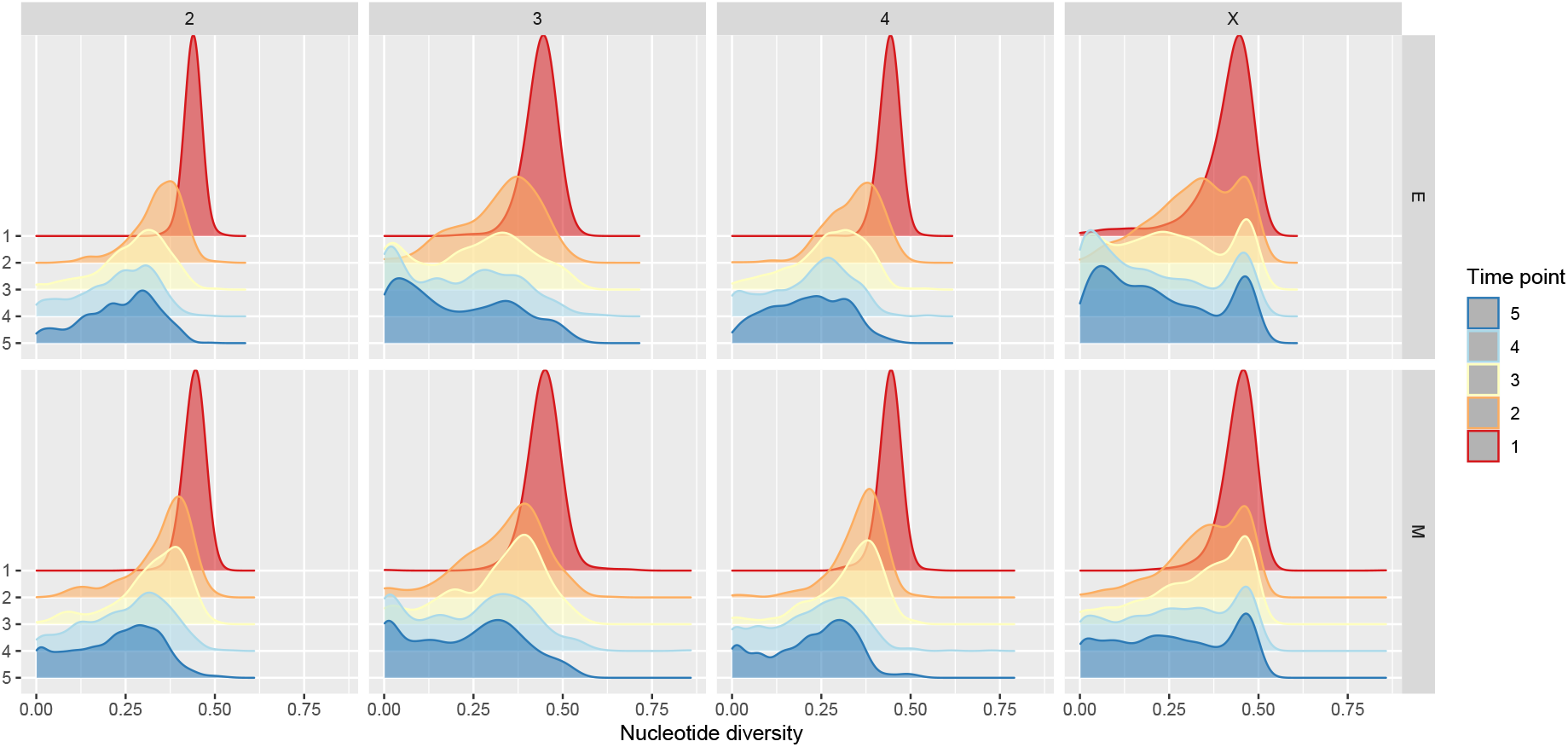
Relative *π* densities in M and E lines. Estimates were computed with Grenedalf (Czech and Exposito-Alonso, 2021). Rows correspond to the two different treatments and columns to chromosomes. Each individual plot has five densities coloured in as per side legend that correspond to one of the five time points.

Tajima’s D estimates throughout the genome are typically greater than 0 but show substantial variation amongst windows (**fig. S9**). These were also computed for 250k SNP windows as for *π*. This result suggests that there is a lack of rare alleles in our dataset which is perhaps unsurprising in a pool-seq experiment. One region worthy of note is the centre of the X chromosome where Tajima’s D is consistently elevated in comparison to surrounding stretches (**fig. 3**). The pattern is present in both E and M populations throughout the whole experiment.

**Figure 3:**
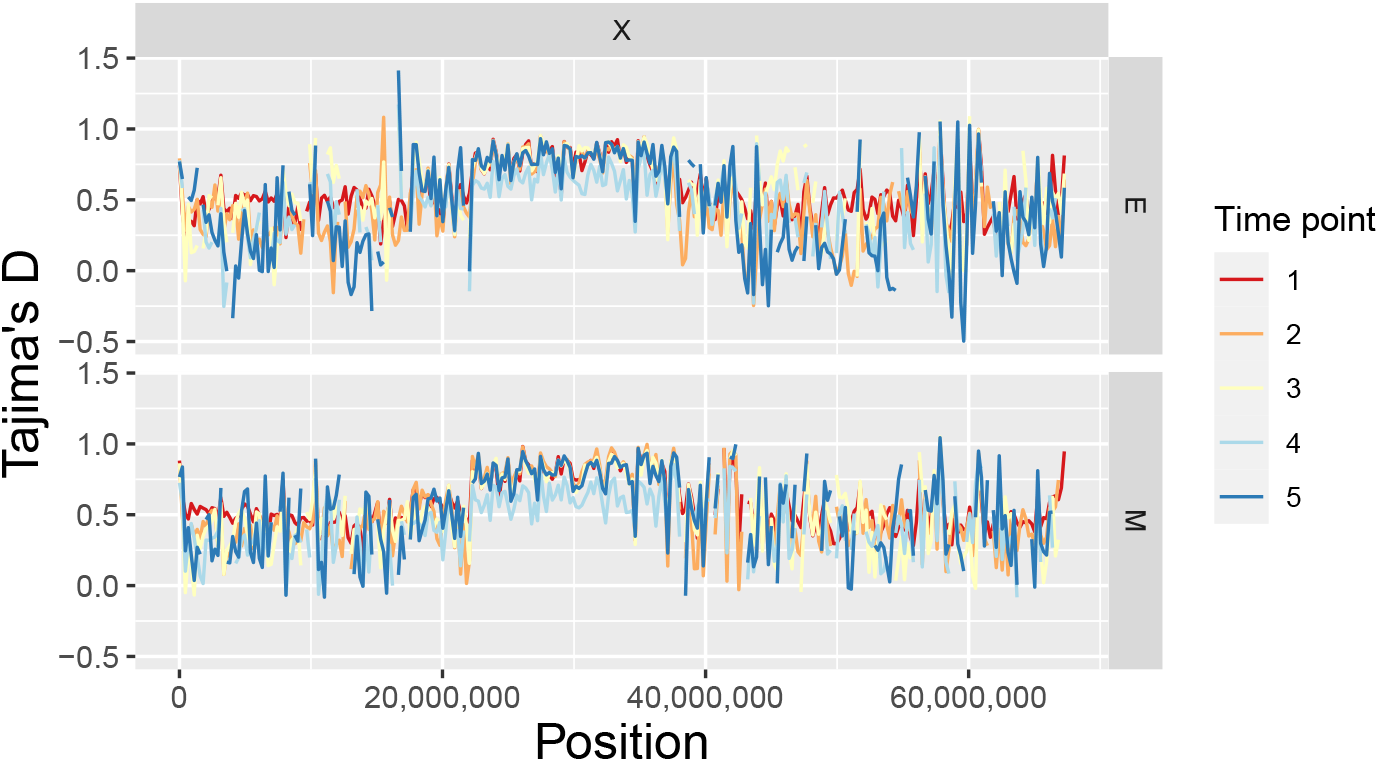
Tajima’s D estimates along the X chromosome for E and M lines. Rows correspond to the two different treatments. Estimates were calculated in 250k SNP windows with Grenedalf (Czech and Exposito-Alonso, 2021). Lines are coloured per time point according to the side legend.

### 2.2 Estimating the effective population size

Estimating the effective population size in windows across the genome should shed light on how fast selection and drift act together to cause allele frequencies to change. Using an estimator that relies on frequencies changing between any two time points (Jóńas et al., 2016), we looked for differences in *N*_*e*_ between chromosomes and treatments. Note that this approach uses data on any polymorphic sites and computes an estimate of variance *N*_*e*_ which does not use any information from fixed loci. Previous results using molecular marker-based estimators, suggest that *N*_*e*_ is similar between lines, ranging from 141.2 (s.d. 27.4) to 110.5 (s.d. 19.2) for M and E, respectively (Snook et al., 2009). Our results confirm this: genome-wide median estimates considering allele frequency changes between time point 1 and 5 are 145 for M populations and 150 for E lines (**table 1**).

**Table 1:** Median genome-wide *N*_*e*_ estimates for M and E lines at different time point intervals. Medians were calculated using 2k SNP window estimates from all of the four experimental replicates. ‘Overall’ corresponds to *N*_*e*_ estimates based on allele frequency changes between the first and last time point. The total number of windows considered in each replicate is found in brackets.

| Time interval | Median - M | Median - E |
| --- | --- | --- |
| Overall | 144.6 (n = 134) | 149.6 (n = 258) |
| T1T2 | 100.6 (n = 292) | 90.7 (n = 360) |
| T2T3 | 67.1 (n = 307) | 110 (n = 288) |
| T3T4 | 66.5 (n = 167) | 120.3 (n = 139) |
| T4T5 | 134.2 (n = 140) | 140.1 (n = 140) |

In E populations, *N*_*e*_ drops most during the first 20 to 60 generations implying that selection is strongest then. *N*_*e*_ starts to recover from time point 3 onwards reaching ≈ 140 at the end of the experiment. Such a result is only possible because variance *N*_*e*_ is estimated from observed temporal shifts in allele frequencies. This is not caused by new mutations as we assume that adaptation occurs from standing genetic variation throughout this study. This same pattern is not found in M lines, where *N*_*e*_ estimates suffer a continuous and strong reduction until time point 4 (from 101 at generation ∼21 to 66 at generation ∼161), after which neutral levels are nearly recovered **table 1**. Similar patterns are recovered when using only intergenic variants to estimate *N*_*e*_ (**table S5**). Since finding that *N*_*e*_ recovers towards the end of the experiment, we compared estimates between the first and last time point. These are significantly different (Mann-Whitney U test *p*-value = 4.4 × 10^−7^), suggesting that (i) selection acting at the start might be causing a significant reduction in *N*_*e*_, and (ii) selection during the last intervals of the experiment is less effective in altering allele frequencies, allowing *N*_*e*_ to recover.

Reduced *N*_*e*_ during the first half of the experiment - time points 1 to 2 - might indicate a strong selective response as E populations reach the new phenotypic optimum very swiftly. In contrast, selection under monogamy seems to act less effectively, causing low *N*_*e*_ until the third quarter of the experiment.

Interestingly, the first interval is marked by low *N*_*e*_ on the X chromosome especially in E populations (**fig. 4**, panel (b)). We investigated whether the estimates on the X differed from those on the autosomes and this difference is statistically significant (**fig. 4**, panel (a), Mann-Whitney U test *p*-value = 9.4 × 10^−6^). Thus, the X chromosome shows a fast adaptive response in E lines from the onset of selection. This result is not as clearly replicated in M populations where *N*_*e*_ estimates on the X are more similar to those on chromosomes 3 and 4 from T1 to T4 (Mann-Whitney U test *p*-value = 0.051).

**Figure 4:**
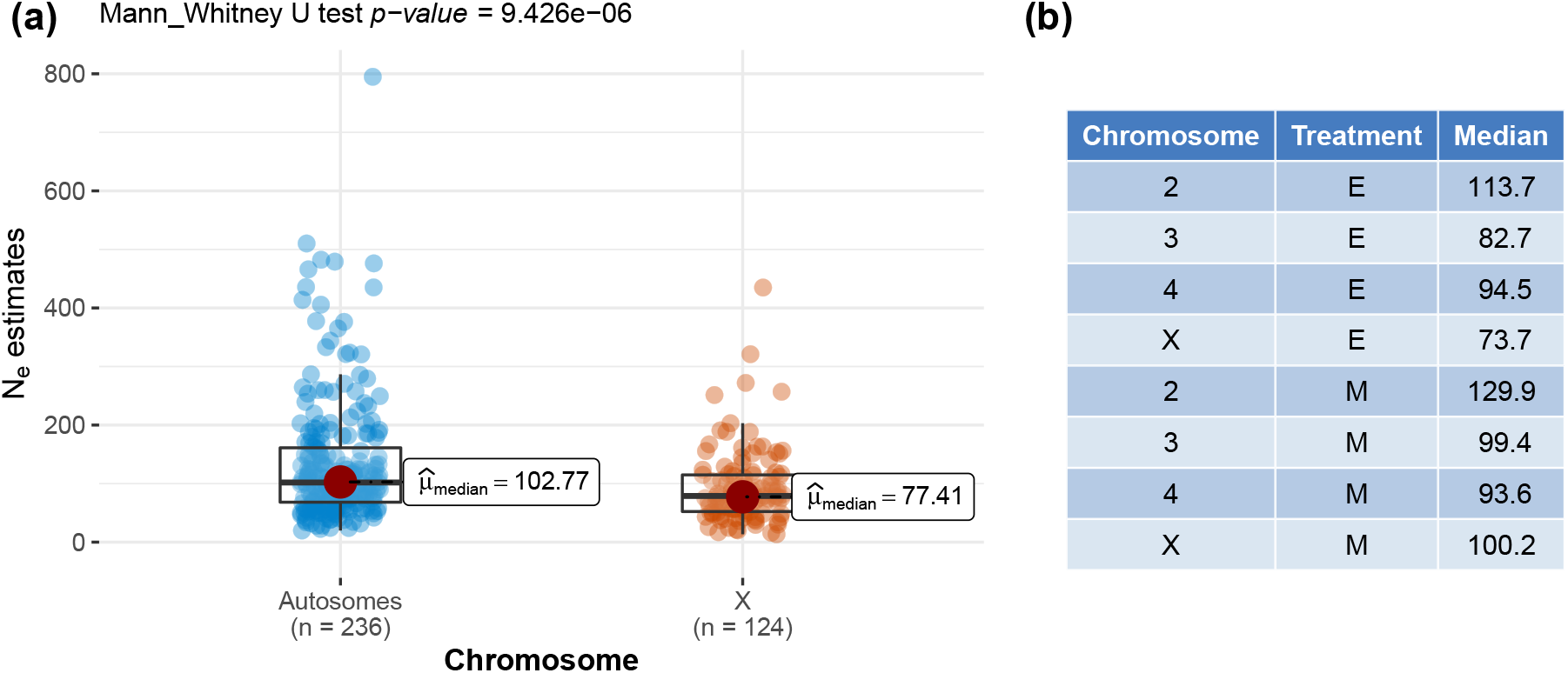
*N*_*e*_ estimate comparison between autosomes and the X chromosome during the first 20 to 60 generations (T1 to T2). Panel (a) shows boxplots of *N*_*e*_ window estimates for the two categories - autosomes and X - in E populations (Mann-Whitney U test very significant). Twelve outliers were removed from (a) for visualisation purposes. Panel (b) shows median estimates per chromosome within this first time point interval.

### 2.3 Estimating selection

We performed a genome scan of the time-series data across all time points using Bait-ER (Barata et al., 2020). The signal of selection is substantially higher in E versus M lines (**fig. 5**), which suggests that selection is indeed stronger under elevated polyandry. In total, 350 (0.9% of all sites in the time-series) and 770 (1.5%) SNPs were statistically significant (at a threshold of *log*(99)) for M and E lines, respectively. If considering the first 3 time points alone, M lines had 570 (0.6%) significant SNPs whilst E had 1591 (1.5%). Regardless of whether you consider the complete time-series or a shorter dataset with the first 3 time point only, E populations have a similar percentage of sites - 1.5% - that are considered to be under selection. They consistently show approximately double the number of loci with evidence of selection than the M lines.

**Figure 5:**
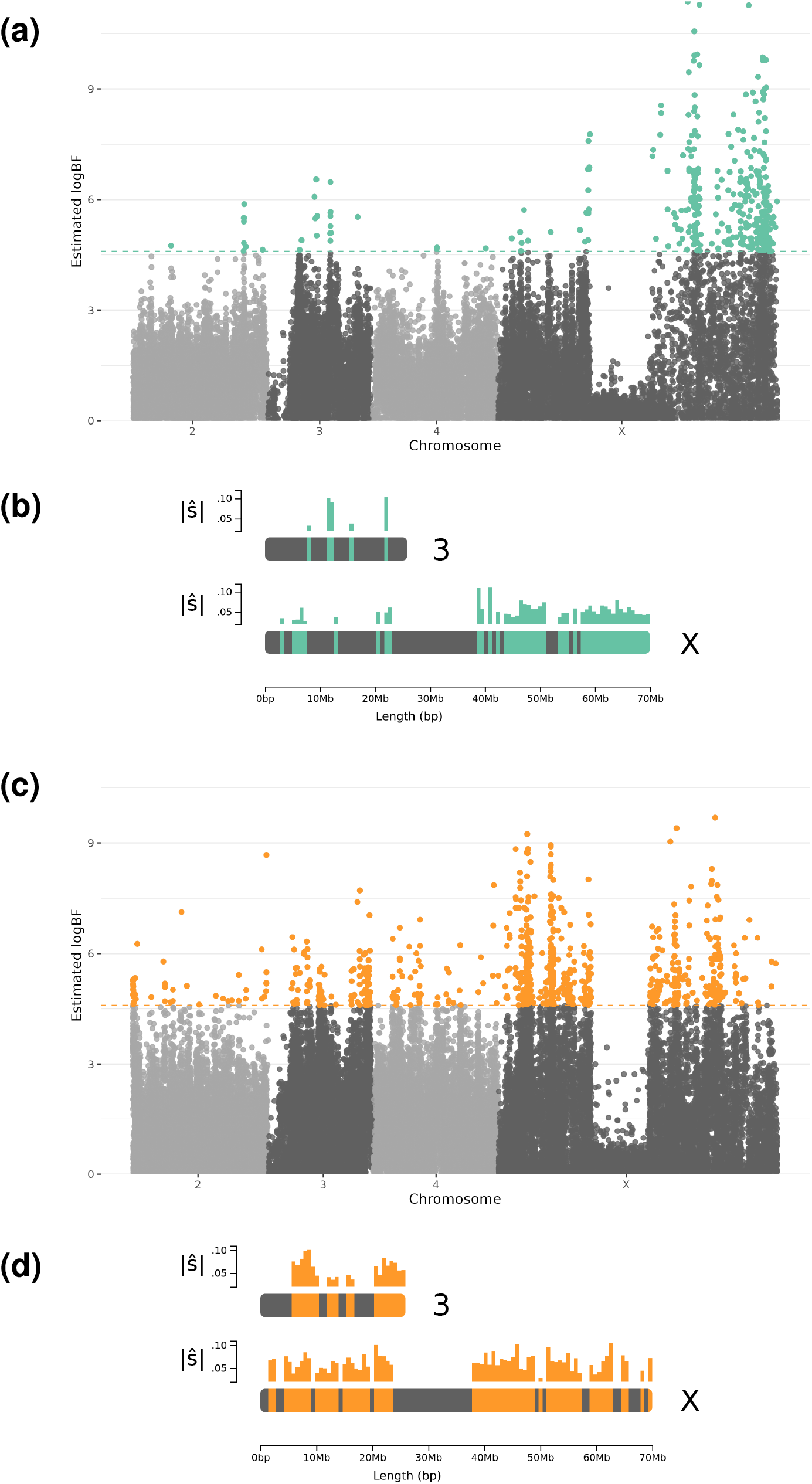
Genome scan for signatures of adaptation throughout the genome for M (top) and E (bottom) lines. (a) and (c) are manhattan plots of Bait-ER (Barata et al., 2020) logBF for each allele frequency trajectory. Statistically significant SNPs are coloured in green (M, top) or orange (E, bottom). Dashed lines correspond to a threshold of *log* (99) ≈ 4.6. (b) and (d) are diagrams of chromosomes 3 (top) and X (bottom) that illustrate which regions of each chromosome harboured the most number of significant hits. Average estimated selection coefficients (| ŝ |) for each interval can be found above each diagram as a bar plot. Data excludes chromosome 5.

When comparing the different chromosomes, it is clear that there are far more significant peaks along the X chromosome in comparison to autosomes in both treatments (M: 322; E:563). Of those 322 top X candidates in M lines, 309 (96%) were located in intergenic regions (versus 50.8% in E). This is a surprising result given that half of the variants called on the X can be found within intergenic regions. Significant trajectories on chromosomes 2 and 4 were never seen for more than 60 SNPs (2: 9 and 43; 4: 3 and 54; for M and E, respectively). In addition, evidence for selection on the 3^rd^ chromosome is also markedly elevated in E populations where there are 110 significant SNPs versus only 16 in M (**fig. 5**). Taken together, these results suggest that whilst selection is stronger under elevated polyandry, the X chromosome is also responsible for adaptation to a strict monogamy regime.

In E lines, 417 (54.2%) of those statistically significant variants were found within genes, whereas only 35 SNPs (10%) were mapped within genes in M lines. This difference is striking given that approximately the same number of significant variants were located in intergenic regions (E: 353; M: 315). A large proportion of the E line top variants that were found in genes locate to the X chromosome (426, 72.3%). Fourteen genes in E populations have 5 or more significant SNPs (up to 23; **table S7**). Selection coefficients for individual trajectories as estimated with Bait-ER ranged from 0.03 to 0.11. Genes with the highest number of top SNPs were found on the X and third chromosomes. Of these, the second gene with the most top variants (11) codes for hemicentin-1 (NCBI: 4813557; FlyBase: FBgn0076932) with an ortholog in *Drosophila melanogaster* - neuromusculin which is a protein that is expressed in the muscle system as well as the peripheral nervous system. Allele frequency trajectories for these top SNPs typically start at relatively high frequency and fix within two or three time points (**fig. 6**).

**Figure 6:**
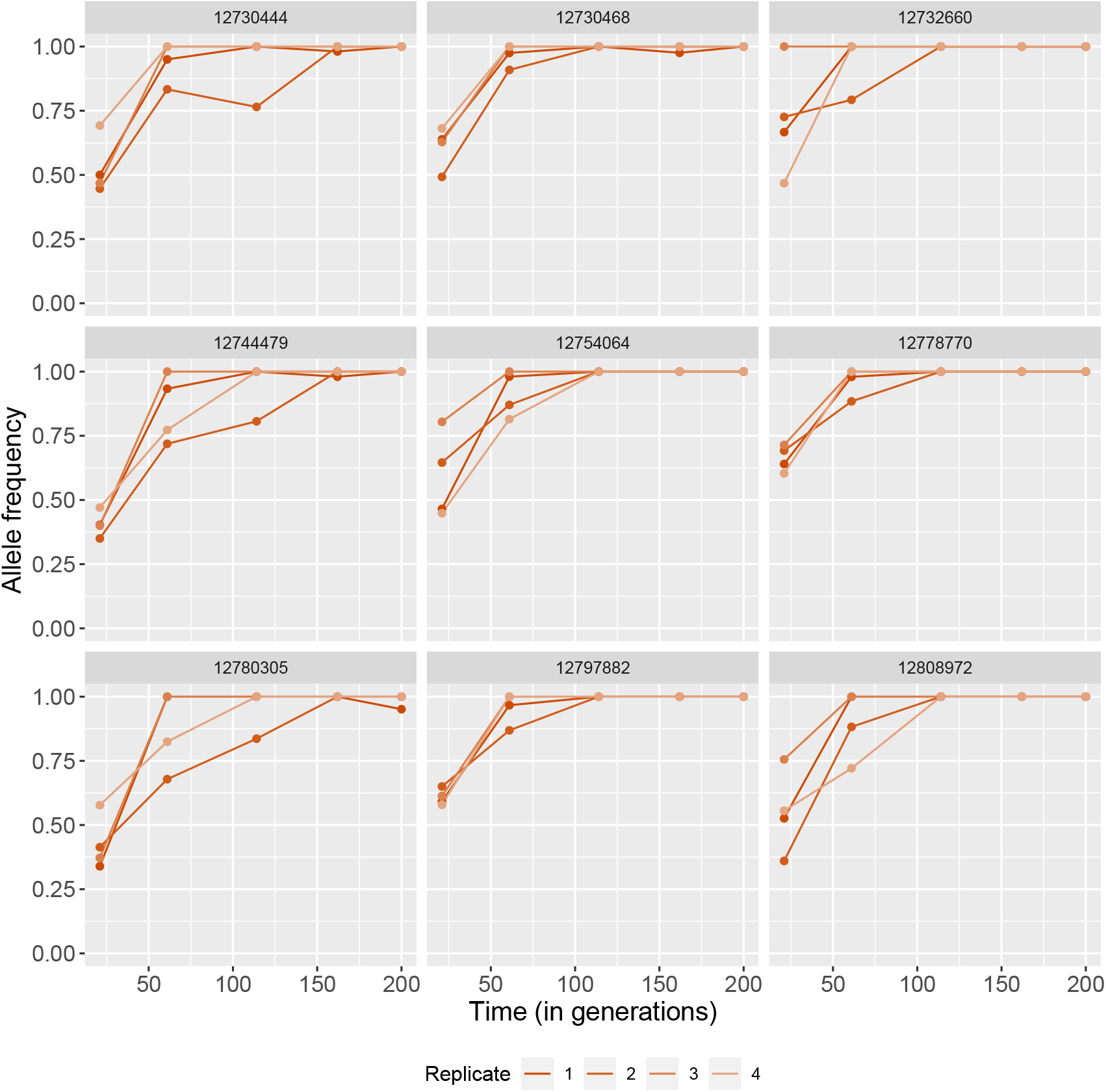
Nine (out of 11) allele frequency trajectories of significant SNPs located on a single gene coding for neuromusculin. Each replicate is coloured differently as per bottom legend. Individual variant coordinates can be found at the top of each graph.

Wiberg et al. (2021) previously identified 480 variants as having a significant allele frequency differences between M and E replicates at time point 3. We determined which genes these top SNPs were located in as well as any genes in the vicinity of intergenic top SNPs. We then compared these genes with those found significant in our genome scan. There were 21 genes in common between the two studies (**table S6**). These include genes involved in neural and muscle development, as well as other biological regulation processes.

## 3 Discussion

Sexual selection can cause substantial divergence between populations and is even believed to be involved in speciation (Seehausen et al., 2014; Janicke et al., 2018). It has been repeatedly implicated in altered ratios of genetic diversity between sex chromosomes and autosomes (e.g. Corl and Ellegren, 2012). Here, we used an E&R experimental design in *D. pseudoobscura* to help elucidate the process of adaptation when the strength of sexual selection is altered in the short-term. We altered the intensity of sexual selection by reducing it in monogamous populations (M) or elevating it in a polyandrous regime (E). Signals of selection were strongest in E populations where sexual selection is elevated. This response is accompanied by a reduction in nucleotide diversity and more alleles becoming fixed as time progresses. In addition, *N*_*e*_ estimates suffer a reduction as populations adapt, but recover towards neutral levels.

While M lines should exhibit relaxed selection since competition for mates is eliminated, E lines are likely to be subject to elevated selection. Increasing the number of males a single female is housed with should cause sexual selection to be stronger. The elevated polyandry regime thus results from the observation that *D. pseudoobscura* are naturally polyandrous and each female has been found to mate with 2 to 3 males within its lifetime (Dobzhansky and Pavlovsky, 1967). Therefore, housing a female with 6 males instead should increase intrasexual competition. One would predict that increased promiscuity would facilitate the evolution of traits involved in mating or fertilisation success and perhaps pre-zygotic isolation mechanisms between the two treatments. Our *D. pseudoobscura* populations show a lack of assortative mating either between treatments or within lines (Debelle et al., 2016). A possible explanation for this might be that male-male competition overcomes the coevolution of female preference in this experiment. This observation could help generate expectations regarding some of the genomic response to selection. Under monogamy, competition for mates is eliminated. In a system where female preference is overshadowed by competition amongst males, sexually antagonistic selection is reduced to a minimum in M lines. If sexual conflict is promptly resolved, the adaptive response to enforced monogamy should be dominated by sexually antagonistic variation being drastically reduced.

Adaptation to an altered mating system will shape patterns of genetic variation in somewhat unpredictable ways. Understanding how allele frequency changes occur within given haplotype structures is instrumental to finding putative targets of selection. Is the signal of adaptation to sexual selection even throughout the genome and consistent across time? Our approach to understanding the adaptive process relied on taking snapshots of the replicates at several time points throughout the 200 generation experiment. These snapshots were allele frequencies estimated from a pool-seq dataset of each of the four replicate populations. Our *two-mapper two-variant caller* approach ensures that only high quality SNPs are present in both the full time-series and the two time point datasets. In particular, selection scan results are based on a time-series that is comprised of SNPs that were polymorphic at the first time sampling point. This causes our results to be focused on polymorphisms with the most adaptive potential, since most would have possibly overcome the counteracting effects of drift over the first 20 generations of selection. In small populations such as these, drift will cause alleles to shift such that most low frequency polymorphisms will be lost within a few generations.

Taken together, our results support the hypothesis that strong sexual selection in E lines causes a substantial adaptive response. Not only did alleles become fixed more promptly in E populations (**fig. 1**), but also nucleotide diversity was depleted faster (**fig. 2**). In addition, our genome scan showed more than double the number of target candidates in E versus M. These top SNPs were found mostly across the X and the 3^rd^ chromosomes (**fig. 5**). These results are consistent with the findings by Wiberg et al. (2021) whose SNPs showing significantly consistent allele frequency differences between E and M clustered along chromosomes 3 and X. The selection signatures we find are more pervasive in comparison to Wiberg et al.’s “islands” of differentiation. This is perhaps suggesting that LD has a substantial impact in our genome scan. In addition, effective population size estimates are consistent with a swift response to selection from the start of the experiment, especially in E. *N*_*e*_ estimated from allele frequency changes between time points 1 and 2 indicate that E lines suffer a more drastic reduction from the onset of selection.

At the end of the first half of the experiment - between time point 2 and 3 - E populations showed far more fixations than M lines (**fig. 1**). We hypothesised that this could result in a more marked response to selection in E which could, in turn, manifest as a reduction in *N*_*e*_ within that time interval. Interestingly, the pattern is reversed when comparing *N*_*e*_ between the two treatments: M lines have a much lower overall *N*_*e*_ on average than E. This result is statistically significant (M versus E at T2 to T3 Mann-Whitney U test p-value = 8 × 10^−13^). This could mean that, despite more overall fixed loci, variance-*N*_*e*_ at the end of this first half indicates a more substantial reduction amongst M populations. A pattern such as this might be caused by one of two things. First, drift could be stronger in M overall resulting in drift variance that is picked up by the *N*_*e*_ estimator. Other monogamous regimes have been shown to result in lower overall *N*_*e*_ which augments the extent of drift (Wigby and Chapman, 2004). *N*_*e*_ in monogamous populations of *Drosophila melanogaster* lines was found to be only 16.2% smaller than that in polyandrous lines (Rice and Holland, 2005). The prediction that *N*_*e*_ differs between the treatments was tested previously in our lines and the difference was found not to be significant (Snook et al., 2009). Secondly, our estimates could be biased if selection affects most allele frequency changes. This effect should dissipate if one would estimate *N*_*e*_ with sites that are evolving neutrally. We tried to overcome this by computing *N*_*e*_ using intergenic SNPs alone. General trends remained unchanged with similar median *N*_*e*_ for M and E between T2 and T3 as that using the complete dataset (M: 68.2 and E: 111.9; **table S5**).

As costs of promiscuity arise from different mating frequency optima between the sexes, monogamy lines should show signs of resolved sexual conflict. This could, in turn, result in a more even distribution of nucleotide diversity along the genome, as well as less variance in *N*_*e*_. Monogamous populations do exhibit less response to selection with fewer significant hits across the genome. This finding can be evidence for relaxed sexual selection due to mate competition being eliminated. Those regions showing selection signatures may have harboured a substantial portion of the sexually antagonistic variation. With most phenotypic optima that differed between the sexes having converged by the end of the experiment, such sexually antagonistic loci would have likely been targets of selection. Interestingly, median *N*_*e*_ in M throughout the experiment is severely reduced until time point 4 to values lower than those found in E. An accelerated rate of genetic drift due to monogamy cannot be ruled out here. However, this could, alternatively, be evidence of a delayed response where new phenotypic optima are reached towards the last quarter of the experiment.

Most of the adaptive signal is found on the X chromosome, which is unsurprising given that most genes involved in the response to both intra- and intersexual conflict are usually expected to be X-linked (Gibson et al., 2002; Connallon and Jakubowski, 2009; Mank and Ellegren, 2009). This signal is widespread across the X in E lines. Evidence for a faster-X effect (Charlesworth et al., 1987) is also supported by a low X/A *N*_*e*_ ratio during the first time point interval. Here, *N*_*e*_ estimates on the X are significantly lower than those on the autosomes (**fig. 4**). Indeed, the substitution of favourable mutations on the X appears swifter than on autosomes. Diversity ratios are also predicted to be reduced as a result of hitchhiking favouring recessive beneficial alleles on the X (Betancourt et al., 2004), an effect that should be elevated under polyandry (Pool and Nielsen, 2008). However, reduced recombination on chromosome X favours the increase in frequency of whole haplotypes. This could be causing the widespread signal that our genome scan is producing. Determining the extent of linkage disequilibrium in this experiment is difficult as there is no data on the haplotypes present in the founder populations in our pool-seq approach. Nevertheless, signatures of adaptation emerging from examining the X chromosome are still compelling.

Our results suggest that the predicted 3*/*4 reduction in *N*_*e*_ on X chromosome is not prevalent throughout the experiment. *N*_*eX*_ is just as high as on the autosomes in M lines regardless of the time point interval in question. Take the example of the T1 and T2 interval, median *N*_*e*_ is very similar for chromosomes 3, 4 and X (**fig. 4**). A similar trend is observed in E lines where *N*_*eX*_ is only found to be the lowest in comparison to any autosome between T1 and T2. If considering any other interval, *N*_*e*_ on the X is as high at that calculated for the autosomes. This is a striking result as all theoretical studies predict a lower *N*_*eX*_ (Pool and Nielsen, 2008; Betancourt et al., 2004), especially under polyandry. One possible explanation might be that balancing selection could be maintaining higher levels of polymorphism which could increase estimates of variance-*N*_*eX*_. In such cases, pervasive balancing selection might arise if a large proportion of this variation is antagonistic between the sexes. Alternatively, most adaptive mutations could be partially dominant, lessening any faster-X effects.

As far as the X chromosome is concerned, the response found on the two chromosome arms differs between the treatments. M lines had a more pronounced response towards the distal end of the chromosome, whereas E lines’ signal was more evenly distributed along the chromosome. This suggests that the genetic basis of adaptation to either elevated or relaxed sexual selection on the X is distinct. This pattern would not be detected in most studies which simply compare divergence in allele frequencies between any two lines. The distal end of the X chromosome (chromosome arm XR in previous *D. pseudoobscura* assemblies) is equivalent to Muller element D in the *D. melanogaster* genome. This chromosome arm is known to have fused with the ancestral X chromosome to form the ”neo-X”. This could indicate that most of the sexually antagonistic variation is found on chromosome arm XR. The signatures found could be a signal of resolved sexual conflict. The ancestral X arm (XL) may have had enough evolutionary time to resolve any intragenomic conflict through mutation and recombination prior to the fusion. Moreover, both treatments exhibited a marked valley of signals of selection (**fig. 5**) in the centre of the X. This was coupled with a positive and elevated Tajima’s D within the same region (**fig. 3**). Such a pattern suggests that, in spite of evidence for positive selection on both chromosome arms, the centromere region could be under balancing selection. This could perhaps be the result of sex-specific allele differentiation on the X between the two sexes. The centre of the X also contains some of the highest coverage regions across the genome (**fig. S10**). This indicates that it might be a highly repetitive portion of the chromosome. Phased data from long read sequencing technology would be necessary to resolve this issue.

One other interesting finding is that populations recovered to neutral levels of *N*_*e*_ towards the middle or end of the experiment for E and M lines, respectively. Here, drift variance caused by any allele frequency changes that match neutral expectations might indicate the end of an initial strong selection phase. It is possible that phenotypic optima are reached at that point and allele frequencies might even plateau. In other words, directional selection becomes less effective towards the end of the experiment as populations reach the new phenotypic optima. This is as if positive selection is relaxed despite no changes to the selective environment. Selection coefficients are reduced and other modes of selection may arise. Again, balancing selection may now be present and act to maintain some genetic diversity. Any remaining genetic variation at the end behaved in such a way that allele frequency variance matched drift expectations. This is independent from any previous allele fixations that might have occurred as a result of directional selection. While more in-depth studies of the genetic basis of sexual conflict would be required to elucidate this matter, our findings support the potential that it has largely been resolved.

Wiberg et al. (2021) found a cluster of top SNPs on chromosome 3 that showed significant differentiation between M and E lines. High levels of nucleotide diversity observed at the start of the experiment (**fig. S8**) make chromosome 3 a good candidate for harbouring selection targets. This would facilitate adaptation due to increased fitness variance amongst individuals in the population. The region at the end of the chromosome was identified by Wiberg et al. as also showing a steep rate of decay in LD. This suggests that this peak region exhibits high recombination which is unexpected given that telomeres are typically low recombination regions. Our study confirms these results. There is evidence for positive directional selection within this region. Interestingly, the signal seems to be caused solely by directional selection in E lines. Increased recombination at the end of chromosome 3 relative to neighbouring areas could have contributed to the slightly elevated nucleotide diversity (**fig. S8**).

In summary, we showed that the response to an altered mating system in populations of *D. pseudoobscura* is found mostly to the X chromosome, but also on chromosome 3. This is consistent with previous work that focused on comparing E and M lines at time point 4 in our analysis (Wiberg et al., 2021). Selection signal is strongest when mate competition is strongest due to elevated polyandry. Such a pattern indicates that most allele frequency changes observed were in fact caused by elevated sexual selection and not solely adaptation to lab conditions. Our study showed the power of investigating allele frequency trajectories and their usefulness when estimating selection parameters and the effective population size.

## 4 Material and Methods

### 4.1 Experimental setup

The experiment was established in 2002 and lasted for approximately 200 generations (Snook et al., 2005). The ancestral population was established from 50 wild-caught females collected in Arizona, USA. The selection experiment was set up after four generations of “common-garden” laboratory evolution. Each of the two treatment lines (M and E) was replicated four times. For each selection regime, recently eclosed offspring were collected and combined given the appropriate sex ratio at every generation. All experimental populations were kept with standard food media and added live yeast at 22°C on a 12-L:12-D light cycle. For a more detailed description, see Snook et al. (2005) and Crudgington et al. (2005).

Both selection regimes were established based on the observation that *D. pseudoobscura* females are known to carry sperm from two males at any given time in the wild (Cobbs, 1977). Therefore, for each E treatment group, two M line groups were established. For each replicate population, 80 and 40 groups of flies remained after culling in M and E treatment lines, respectively. Consequently, we expect no differences in the potential for adaptation between treatments due to reduced effective population size.

### 4.2 Sequencing

The time-series dataset consists of 5 time points and it includes all four replicates for each selection regime. These time points are fairly evenly distributed throughout the study. The experimental setup was such that the replicates were established in a staggered fashion. Therefore, fly sampling did not occur at the same for all replicate populations: time point 1 was sampled at generations 21 and 22, time point 2 between 59-63, time point 3 between 112-116, time point 4 ranged between 160-164, and time point 5 at generation 200. For more details on the generation at which each replicate was sampled and, thus, sequenced, see **table S1**. Samples at time point 4 were sequenced as part of Wiberg et al. (2021).

All fly samples were stored at -80°C immediately after collection in the Department of Animal and Plant Sciences at the University of Sheffield. The samples were then collected and kept at -80°C storage in the Centre for Biological Diversity at the University of St Andrews up until DNA extraction.

For each DNA sample, 40 female flies were pooled from the frozen stocks. Females were sexed and collected approximately at time of emergence, thus, we assume these to be virgin. DNA extraction was performed using a DNeasy Blood & Tissue Kit (QIAGEN) for 20 individuals. Firstly, flies were homogenised at room temperature using a Bullet Blender homogeniser with zirconium beads. After adding Proteinase K, all samples were left to incubate overnight at 56°C. The step which involves adding buffer AW1 was repeated, and the elution with buffer AE (150 μL) was also repeated to maximise DNA yield. At the end of the extraction protocol, the two 20-female samples were combined to make up a pool of 40 females and stored at -20°C. DNA concentration increased as time progresses, with time point 1 at 34.9 *ng /*μ*L*, time point 2 at 45.1 *ng /*μ*L*, time point 3 at 45.4 *ng /*μ*L*, and time point 5 at 50.8 *ng /*μ*L*. This is consistent with more recent DNA samples being better preserved.

DNA sequencing was carried out at Novogene (Hong Kong) using an Illumina HiSeq X Ten platform. The library preparation protocol resulted in a 350bp insert DNA library. For each sample, there is a set of raw paired-end reads all 150bp long.

### 4.3 Read mapping

Raw reads were filtered and trimmed using Trimmomatic (version 0.38, Bolger et al., 2014). After trimming, time points 1, 2, 3, and 5 had an average read length of 148 (min = 36, max = 150), whilst time point 4 had shorter reads with an average length of 97.3 (min = 36, max = 100). Trimmed reads were then mapped to both the complete *Drosophila pseudoobscura* genome assembly Dpse 4.0 (FlyBase, GenBank accession GCA 000001765.3, June 2018) and the X chromosome sequence of the UCBerk Dpse 1.0 assembly (UC Berkeley, GenBank accession GCA 004329205.1, March 2019). The former reference was assembled with Illumina (150x) and PacBio (70x) reads, and the latter consists of Oxford Nanopore MinION (40x) long reads.

All paired-end reads were mapped separately using two mappers: bwa mem (version 0.7.17, default parameters, Li, 2013) and novoalign (version 4.00.Pre-20190624, Novocraft Technologies, http://novocraft.com/). Regardless of which mapper was used, over 98% of reads were mapped successfully to the reference genome (see **table S2** for more details). The SAM files produced by the two mappers were re-aligned to around indels using GATK (Genome Analysis Tool Kit, version 3.8.1, Van der Auwera and O’Connor, 2020).

### 4.4 Variant calling and filtering

Variants were then called with both bcftools (mpileup and call functions, version 1.9, Li, 2011; Danecek et al., 2021) and freebayes (version 1.3.3, Garrison and Marth, 2012) (see **table S3** for details on variants called by both callers). Variants called by bcftools were filtered according to the following criteria suggested by Kofler et al. (2016):

1. Minimum mapping quality of 40;
2. Minimum base quality of 30;
3. Minimum allele count of 1/F at the first time point, where F is the total number of founder haplotypes or the sample size, i.e. MAF = 1/40 = 0.025;
4. Remove sites not called by FreeBayes;

In addition, we have filtered out those variants not present in both the bwa mem and novoalign mapped datasets. Such a two mapper approach to producing pool-seq data is rather conservative and was preferred to ensure good quality datasets. On average, novoalign alignments led to fewer called mean SNPs genome-wide: 2,309,226 with bwa mem versus 2,194,721 with novoalign for M lines, and 2,281,034 with bwa mem versus 2,167,341 with novoalign for E populations. A similar trend is observed in the X chromosome alignment (bwa mem 862,881 and 841,935 vs novoalign 807,689 and 787,561 for M and E, respectively). Fewer variants were called as time progressed. On average, a little over a third of variants that were called on autosomes were intergenic (chromosome 2: 34.1%; chromosome 3: 34.1%; chromosome 4: 38.6%). A higher percentage was observed on the X chromosome where 47.5% of variants were intergenic. We also considered strand bias (quantity that measures whether a SNP is just as likely to be found on the forward and the reverse strand; it is calculated as |*f*_*forward*_ − 0.5| where *f*_*forward*_ is the proportion of reads that were mapped to the forward strand at any given site; **fig. S4**) and overall coverage for each analysed locus as a measure of quality (**fig. S5**). After filtering for mapping and base calling quality, as well as retaining any variants called by bcftools and Freebayes in the two alignments produced with bwa mem and novoalign, median sequencing depth is 45x. For this reason, we decided not to filter variants based on coverage any further. Finally, we have only considered biallelic sites and of those only the ones that were found to be polymorphic at the first sampled time point were used for further analyses. This allowed us to describe allele frequency changes throughout the experiment. The total number of SNPs remaining after quality filtering as well as those called in both bwa mem and novoalign alignments for the whole-genome and the X chromosome assemblies can be found in **table S3**. After filtering, SNPs present across all replicates between the five time points (or the first three) were considered for further selection inference. We analysed 38,065 and 51,339 full five time point trajectories in M and E lines, respectively. For *N*_*e*_ estimation, those variants present between any two time points were considered. More details on how these were distributed across time points and chromosomes can be found in **table S4** and **fig. S6**.

To investigate how quickly alleles were fixing throughout the experiment, we calculated experimental fixation rates that correspond to the number of SNPs that become fixed from one time point to the next.

Experimental fixation rates were calculated as follows:

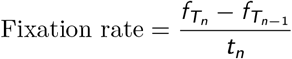

where 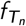 and 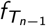 are the number of fixed sites at time point *n* and time point *n* − 1, and *t*_*n*_ is the total number of trajectories being analysed.

### 4.5 Genetic diversity

We used Grenedalf to calculate estimates of nucleotide diversity and Tajima’s D (Czech and Exposito-Alonso, 2021). Grenedalf computes measures of diversity corrected for pool-seq. This includes *θ*_*π*_, hereinafter referred to as *π*. The program follows the approach implemented in PoPoolation (Kofler et al., 2011a) and PoPoolation2 (Kofler et al., 2011b) that accounts for any bias introduced by sampling and sequencing error. Absolute *π* is the sum of estimates for all SNPs within a given window, and relative *π* the average per window. We used sync format files, which include all replicates and time points, as input. We considered sample sizes of 40 individuals and computed *π* as well as Tajima’s D for each replicate at each time point in windows of 250kbp with a 25kbp overlap.

### 4.6 Selection inference and *N*_*e*_ estimation

For estimating the effective population size, *N*_*e*_, we used a moment-based estimator (Jóńas et al., 2016) implemented in the R package poolSeq (Taus et al., 2017). The estimator uses temporal data to investigate any allele frequency changes. It accounts for the effect of pooled sequencing by introducing variance from two sampling events: first, when individuals are sampled from an experimental population for sequencing, and second, due to uneven coverage throughout the genome. The approach models drift variance to obtain a temporal estimator for *N*_*e*_ (also referred to as variance-*N*_*e*_). Estimates were computed for 2k SNP windows with a 10% overlap assuming that populations were sampled according to Jóńas et al.’s plan II, where sampling takes place before reproduction and sampled individuals’ genomes do not contribute to the next generation. Replicate medians are computed first. These are, in turn, used to calculate chromosome-level median estimates. Similarly, each genome-wide *N*_*e*_ estimate is a median of all replicate medians.

Finally, we investigated potential targets of selection using a Bayesian genome scan on the time-series - Bait-ER (Barata et al., 2020). It was designed for E&R experiments as it accounts for added binomial, or beta-binomial, sampling noise from pool-seq. Bait-ER models the evolution of an allele using a Moran model with overlapping generations. It estimates parameters of selection, namely selection coefficients (*σ*), whilst also testing each allele frequency trajectory for selection. The program outputs a Bayes Factor (logBF) per site, which is a ratio of the likelihoods of two alternative models: one where genetic drift is the main driver of allele frequency changes, and another where there is positive selection favouring a particular allele. Similar to Grenedalf, both Bait-ER and poolSeq take sync files as input. Diagrams of chromosomal regions showing significant hits (**fig. 5, b and d**) were produced using the chromoMap R package (Anand and Rodriguez Lopez, 2022).

### 4.7 Gene feature analysis

In order to obtain a complete annotation of gene features for our time-series dataset, we used NCBI’s Remap tool (www.ncbi.nlm.nih.gov/genome/tools/remap). This tools allowed us to perform coordinate remapping between the latest annotated reference genome uploaded to NCBI’s repository (University of California, Irvine, Dpse MV25, accession number GCA 009870125.2) and the two assemblies described in section 4.2. The software outputs gff3 format annotation files which we converted to bed format using BEDOPS’ (Neph et al., 2012) gff2bed tool.

## Abbreviations

AFC: allele frequency change
bp: base-pairs
E&R: Evolve & Resequence
M: monogamy
E: elevated polyandry
pool-seq: pooled sequencing
SNP: single nucleotide polymorphism

## 5 Acknowledgements

This work was supported by the Vienna Science and Technology Fund (WWTF) through project MA16-064. CK received funding from the Royal Society (RG170315) and the Carnegie Trust (RIG007474). MGR and RRS have been supported by NERC (UK) grants NE/I014632/1 and NE/V001566/1. Bioinformatics analyses were performed on the computer cluster at the University of St Andrews Bioinformatics Unit, which is funded by Wellcome Trust ISSF awards 105621/Z/14/Z. Complementary data parsing was carried out with the computational resources provided by the Research/Scientific Computing teams at The James Hutton Institute and the National Institute of Agricultural Botany (NIAB) - UK’s Crop Diversity Bioinformatics HPC, BBSRC grant BB/S019669/1. We are thankful to Paris Veltsos and R. Axel W. Wiberg for useful discussions about the project as well as providing us with the resequencing data they had produced as a result of previous work on this experiment. We are especially grateful to Tanya Sneddon for her help with the DNA extraction process and shipping.

## 6 Data availability

The genomic data generated in this article can be found in the NCBI’s Sequence Read Archive (SRA, https://www.ncbi.nlm.nih.gov/sra). Raw reads were deposited under the BioProject ID PRJNA808747. All scripts used for read mapping, variant calling and filtering, estimating *N*_*e*_ across the genome and inferring selection from standing genetic variation can be found here: https://github.com/carolbarata/dpseudo-n-beyond.

## References

L. Anand and C. M. Rodriguez Lopez. ChromoMap: an R package for interactive visualization of multi-omics data and annotation of chromosomes. BMC Bioinformatics, 23(1):1–9, 2022.

C. Barata, R. Borges, and C. Kosiol. Bait-ER: A Bayesian method to detect targets of selection in evolve-and-resequence experiments. bioRxiv, 2020. URL https://doi.org/10.1101/2020.12.15.422880.

N. Barghi, R. Tobler, V. Nolte, and C. Schlötterer. Drosophila simulans: A Species with Improved Resolution in Evolve and Resequence Studies. G3: Genes—Genomes—Genetics, 7(7):2337–2343, 2017.

A. J. Betancourt, Y. Kim, and H. A. Orr. A pseudohitchhiking model of X vs. autosomal diversity. Genetics, 168(4):2261–2269, 2004.

A. M. Bolger, M. Lohse, and B. Usadel. Trimmomatic: A flexible trimmer for Illumina sequence data. Bioinformatics, 30(15):2114–2120, 2014.

M. K. Burke, J. P. Dunham, P. Shahrestani, K. R. Thornton, M. R. Rose, and A. D. Long. Genome-wide analysis of a long-term evolution experiment with Drosophila. Nature, 467(7315):587–590, 2010.

T. Chapman, L. F. Liddle, J. M. Kalb, M. F. Wolfner, and L. Partridge. Cost of mating in Drosophila melanogaster females is mediated by male accessory gland products. Nature, 373(6511):241–244, 1995.

B. Charlesworth, J. A. Coyne, and N. H. Barton. The Relative Rates of Evolution of Sex Chromosomes and Autosomes. The American Naturalist, 130(1):113–146, 1987.

S. F. Chenoweth and M. W. Blows. Contrasting Mutual Sexual Selection on Homologous Signal Traits in Drosophila serrata. The American Naturalist, 165(2):281–289, 2005.

S. F. Chenoweth, N. C. Appleton, S. L. Allen, and H. D. Rundle. Genomic Evidence that Sexual Selection Impedes Adaptation to a Novel Environment. Current Biology, 25(14):1860–1866, 2015.

A. K. Chippindale, J. R. Gibson, and W. R. Rice. Negative genetic correlation for adult fitness between sexes reveals ontogenetic conflict in Drosophila. Proceedings of the National Academy of Sciences of the United States of America, 98(4):1671–1675, 2001.

G. Cobbs. Multiple Insemination and Male Sexual Selection in Natural Populations of Drosophila pseudoobscura. The American Naturalist, 111(980):641–656, 1977.

T. Connallon and E. Jakubowski. Association between sex ratio distortion and sexually antagonistic fitness consequences of female choice. Evolution, 63(8):2179–2183, 2009.

A. Corl and H. Ellegren. The genomic signature of sexual selection in the genetic diversity of the sex chromosomes and autosomes. Evolution, 66(7):2138–2149, 2012.

H. S. Crudgington, A. P. Beckerman, L. Brüstle, K. Green, and R. R. Snook. Experimental Removal and Elevation of Sexual Selection: Does Sexual Selection Generate Manipulative Males and Resistant Females? The American Naturalist, 165(S5):S72–S87, 2005.

H. S. Crudgington, S. Fellows, N. S. Badcock, and R. R. Snook. Experimental manipulation of sexual selection promotes greater male mating capacity but does not alter sperm investment. Evolution, 63(4): 926–938, 2009.

H. S. Crudgington, S. Fellows, and R. R. Snook. Increased opportunity for sexual conflict promotes harmful males with elevated courtship frequencies. Journal of Evolutionary Biology, 23(2):440–446, 2010.

L. Czech and M. Exposito-Alonso. Grenedalf: Genome Analyses of Differential Allele Frequencies. Manuscript in preparation, 2021.

P. Danecek, J. K. Bonfield, J. Liddle, J. Marshall, V. Ohan, M. O. Pollard, A. Whitwham, T. Keane, S. A. McCarthy, R. M. Davies, and H. Li. Twelve years of SAMtools and BCFtools. GigaScience, 10(2):1–4, 2021.

A. Debelle, M. G. Ritchie, and R. R. Snook. Evolution of divergent female mating preference in response to experimental sexual selection. Evolution, 68(9):2524–2533, 2014.

A. Debelle, M. G. Ritchie, and R. R. Snook. Sexual selection and assortative mating: An experimental test. Journal of Evolutionary Biology, 29(7):1307–1316, 2016.

A. Debelle, A. Courtiol, M. G. Ritchie, and R. R. Snook. Mate choice intensifies motor signalling in Drosophila. Animal Behaviour, 133:169–187, 2017.

T. Dobzhansky and O. Pavlovsky. Repeated Mating and Sperm Mixing in Drosophila pseudoobscura. The American Naturalist, 101(922):527–533, 1967.

J. D. Fry. The Genomic Location of Sexually Antagonistic Variation: Some Cautionary Comments. Evolution, 64(5):1510–1516, 2009.

E. Garrison and G. Marth. Haplotype-based variant detection from short-read sequencing. arXiv, 2012. URL http://arxiv.org/abs/1207.3907.

J. R. Gibson, A. K. Chippindale, and W. R. Rice. The X chromosome is a hot spot for sexually antagonistic fitness variation. Proceedings of the Royal Society B: Biological Sciences, 269(1490):499–505, 2002.

P. C. Griffin, S. B. Hangartner, A. Fournier-Level, and A. A. Hoffmann. Genomic trajectories to desiccation resistance: Convergence and divergence among replicate selected Drosophila lines. Genetics, 205(2): 871–890, 2017.

D. Haig. Intragenomic politics. Cytogenetic and Genome Research, 113(1-4):68–74, 2006.

B. Holland and W. R. Rice. Experimental removal of sexual selection reverses intersexual antagonistic coevolution and removes a reproductive load. Proceedings of the National Academy of Sciences, 96(9): 5083–5088, 1999.

B. Hollis, D. Houle, Z. Yan, T. J. Kawecki, and L. Keller. Evolution under monogamy feminizes gene expression in Drosophila melanogaster. Nature Communications, 5:1–5, 2014.

B. Hollis, L. Keller, and T. J. Kawecki. Sexual selection shapes development and maturation rates in Drosophila. Evolution, 71(2):304–314, 2017.

E. Immonen, R. R. Snook, and M. G. Ritchie. Mating system variation drives rapid evolution of the female transcriptome in Drosophila pseudoobscura. Ecology and Evolution, 4(11):2186–2201, 2014.

P. Innocenti and E. H. Morrow. The Sexually Antagonistic Genes of Drosophila melanogaster. PLOS Biology, 8(3):e1000335, 2010.

T. Janicke, M. G. Ritchie, E. H. Morrow, and L. Marie-Orleach. Sexual selection predicts species richness across the animal kingdom. Proceedings of the Royal Society B: Biological Sciences, 285(1878), 2018.

Á. Jóńas, T. Taus, C. Kosiol, C. Schlötterer, and A. Futschik. Estimating the effective population size from temporal allele frequency changes in experimental evolution. Genetics, 204(2):723–735, 2016.

R. Kofler, P. Orozco-terWengel, N. De Maio, R. V. Pandey, V. Nolte, A. Futschik, C. Kosiol, and C. Schlötterer. PoPoolation: A Toolbox for Population Genetic Analysis of Next Generation Sequencing Data from Pooled Individuals. PLOS ONE, 6(1):e15925, 2011a.

R. Kofler, R. V. Pandey, and C. Schlötterer. PoPoolation2: Identifying differentiation between populations using sequencing of pooled DNA samples (Pool-Seq). Bioinformatics, 27(24):3435–3436, 2011b.

R. Kofler, A. M. Langmüller, P. Nouhaud, K. A. Otte, and C. Schlötterer. Suitability of Different Mapping Algorithms for Genome-Wide Polymorphism Scans with Pool-Seq Data. G3: Genes—Genomes—Genetics, 6(November):3507–3515, 2016.

H. Li. A statistical framework for SNP calling, mutation discovery, association mapping and population genetical parameter estimation from sequencing data. Bioinformatics, 27(21):2987–2993, 2011.

H. Li. Aligning sequence reads, clone sequences and assembly contigs with BWA-MEM. arXiv, 2013. URL http://arxiv.org/abs/1303.3997.

J. E. Mank and H. Ellegren. Sex-linkage of sexually antagonistic genes is predicted by female, but not male, effects in birds. Evolution, 63(6):1464–1472, 2009.

S. Neph, M. S. Kuehn, A. P. Reynolds, E. Haugen, R. E. Thurman, A. K. Johnson, E. Rynes, M. T. Maurano, J. Vierstra, S. Thomas, R. Sandstrom, R. Humbert, and J. A. Stamatoyannopoulos. BEDOPS: High-performance genomic feature operations. Bioinformatics, 28(14):1919–1920, 2012.

J. E. Pool and R. Nielsen. The impact of founder events on chromosomal variability in multiply mating species. Molecular Biology and Evolution, 25(8):1728–1736, 2008.

W. R. Rice. Sex chromosomes and the evolution of sexual dimorphism. Evolution, 38(4):735–742, 1984.

W. R. Rice and B. Holland. Experimentally enforced monogamy: inadvertent selection, inbreeding, or evidence for sexually antagonistic coevolution? Evolution, 59(3):682–685, 2005.

F. Ruzicka and T. Connallon. Is the X chromosome a hot spot for sexually antagonistic polymorphisms? Biases in current empirical tests of classical theory: Sexual antagonism on the X chromosome? Proceedings of the Royal Society B: Biological Sciences, 287(1937), 2020.

F. Ruzicka, M. S. Hill, T. M. Pennell, I. Flis, F. C. Ingleby, R. Mott, K. Fowler, E. H. Morrow, and M. Reuter. Genome-wide sexually antagonistic variants reveal long-standing constraints on sexual dimorphism in fruit flies. PLOS Biology, 17(4):e3000244, 2019.

O. Seehausen, R. K. Butlin, I. Keller, C. E. Wagner, J. W. Boughman, P. A. Hohenlohe, C. L. Peichel, G. P. Saetre, C. Bank, Å. Brännström, A. Brelsford, C. S. Clarkson, F. Eroukhmanoff, J. L. Feder, M. C. Fischer, A. D. Foote, A. D. Foote, P. Franchini, C. D. Jiggins, F. C. Jones, A. K. Lindholm, K. Lucek, M. E. Maan, D. A. Marques, S. H. Martin, B. Matthews, J. I. Meier, M. Möst, M. W. Nachman, E. Nonaka, D. J. Rennison, J. Schwarzer, E. T. Watson, A. M. Westram, and A. Widmer. Genomics and the origin of species. Nature Reviews Genetics, 15(3):176–192, 2014.

R. R. Snook, A. Robertson, H. S. Crudgington, and M. G. Ritchie. Experimental manipulation of sexual selection and the evolution of courtship song in Drosophila pseudoobscura. Behavior Genetics, 35(3): 245–255, 2005.

R. R. Snook, L. Brüstle, and J. Slate. A test and review of the role of effective population size on experimental sexual selection patterns. Evolution, 63(7):1923–1933, 2009.

T. Taus, A. Futschik, and C. Schlötterer. Quantifying Selection with Pool-Seq Time Series Data. Molecular Biology and Evolution, 34(11):3023–3034, 2017.

G. A. Van der Auwera and B. D. O’Connor. Genomics in the Cloud: Using Docker, GATK, and WDL in Terra. O’Reilly Media, Sebastopol, CA, USA, 2020. ISBN 9781491975169.

P. Veltsos, Y. Fang, A. R. Cossins, R. R. Snook, and M. G. Ritchie. Mating system manipulation and the evolution of sex-biased gene expression in Drosophila. Nature Communications, 8(1):1–11, 2017.

R. A. W. Wiberg, P. Veltsos, R. R. Snook, and M. G. Ritchie. Experimental evolution supports signatures of sexual selection in genomic divergence. Evolution Letters, 5(3):214–229, 2021.

S. Wigby and T. Chapman. Female resistance to male harm evolves in response to manipulation of sexual conflict. Evolution, 58(5):1028–1037, 2004.

A. E. Wright, M. Fumagalli, C. R. Cooney, N. I. Bloch, F. G. Vieira, S. D. Buechel, N. Kolm, and J. E. Mank. Male-biased gene expression resolves sexual conflict through the evolution of sex-specific genetic architecture. Evolution Letters, 2(2):52–61, 2018.

L. Yun, M. Bayoumi, S. Yang, P. J. Chen, H. D. Rundle, and A. F. Agrawal. Testing for local adaptation in adult male and female fitness among populations evolved under different mate competition regimes. Evolution, 73(8):1604–1616, 2019.

