## Supplementary information for "Selection on the fly: short term adaptation to an altered sexual selection regime in *Drosophila pseudoobscura*"

#### Mapping statistics

| | Generation | | | | |
|---|---|---|---|---|---|
| **Replicate** | **TP 1** | **TP 2** | **TP 3** | **TP 4** | **TP 5** |
| M/E 1 | 21 | 63 | 116 | 164 | 200 |
| M/E 2 | 21 | 62 | 115 | 163 | 200 |
| M/E 3 | 21 | 61 | 114 | 160 | 200 |
| M/E 4 | 22 | 59 | 112 | 160 | 200 |

Table S1: **Description of which generations were sampled at each time point.** Both M and E lines were sampled at the same generation for each corresponding replicate population. TP: time point.

| Treatment | Mapper | Average no. | Average % | Minimum no. (%) | Maximum no. (%) | Assembly |
|---|---|---|---|---|---|---|
| M | bwa | 48,315,592 | 99.0 | 37,798,247 (98.6) | 62,704,196 (98.8) | Whole genome |
| M | novoalign | 48,629,415 | 99.2 | 38,092,806 (98.9) | 61,869,582 (99.2) | Whole genome |
| E | bwa | 52,169,450 | 99.0 | 43,105,534 (98.8) | 65,961,589 (99.2) | Whole genome |
| E | novoalign | 52,568,742 | 99.2 | 43,479,141 (99.1) | 66,521,236 (99.3) | Whole genome |
| M | bwa | 22,269,730 | 98.4 | 17,630,965 (98.1) | 28,450,846 (98.3) | X chromosome |
| M | novoalign | 22,712,923 | 98.8 | 17,883,361 (98.6) | 29,118,289 (98.8) | X chromosome |
| E | bwa | 24,014,102 | 98.4 | 19,722,067 (98.3) | 29,666,606 (98.8) | X chromosome |
| E | novoalign | 24,494,392 | 98.9 | 20,185,805 (98.8) | 30,279,204 (99.0) | X chromosome |

Table S2: **Mapping statistics for both mappers.** This includes average number of mapped reads as well as the average percentage across samples for M and E lines. Data on whole genome and X chromosome level assemblies can be found on this table.

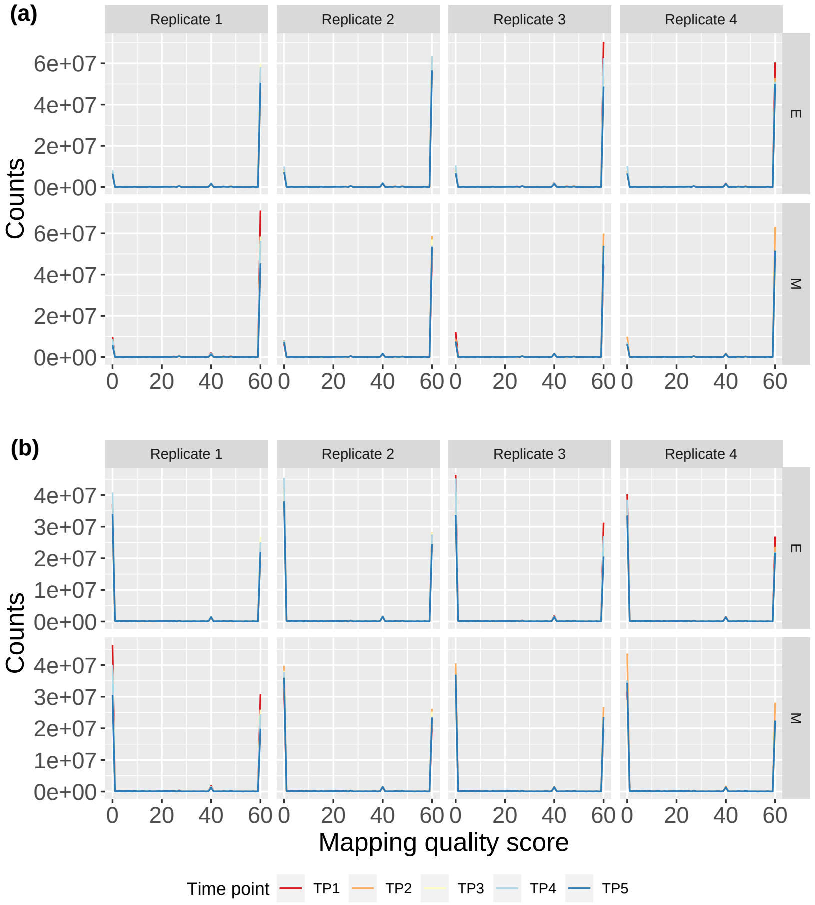


Figure S1: **Mapping quality score distribution at the genome (a) and X chromosome (b) level assemblies.** Rows correspond to the two treatments, E (top) and M (bottom), and columns to the four experimental replicates. Each time point is coloured differently as per legend at the bottom.

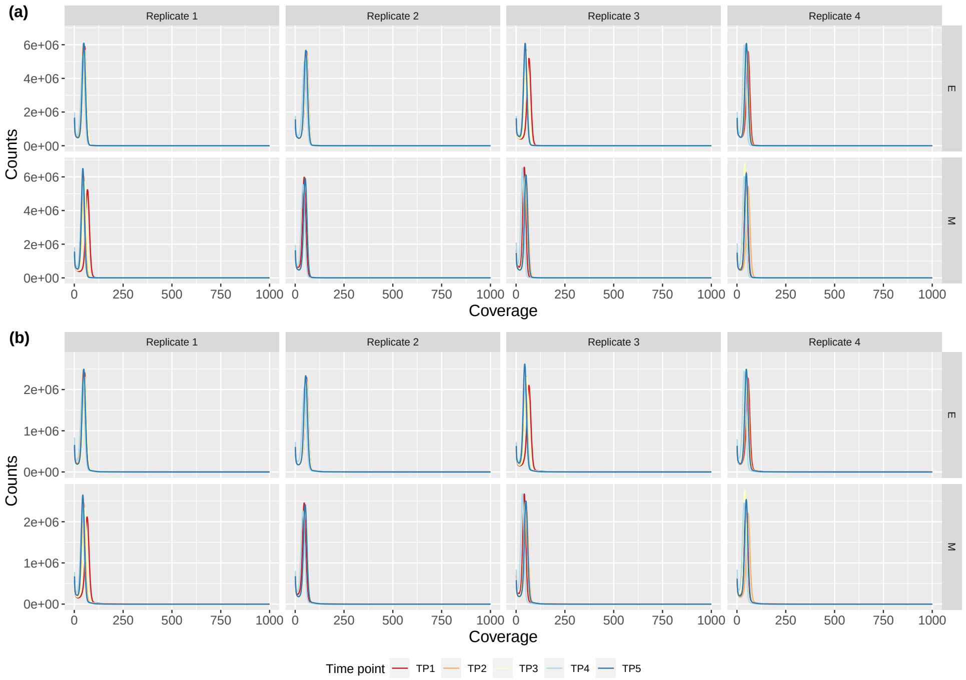


Figure S2: **Read coverage distribution at the genome and X chromosome level assemblies.** Each column corresponds to an different replicate population and each row to either E (top) or M (bottom) lines. The top panel - (a) - shows genome level data and the bottom panel - (b) - X chromosome data. Different time points are coloured according to legend at the bottom of the figure.

### Variant calling and filtering statistics

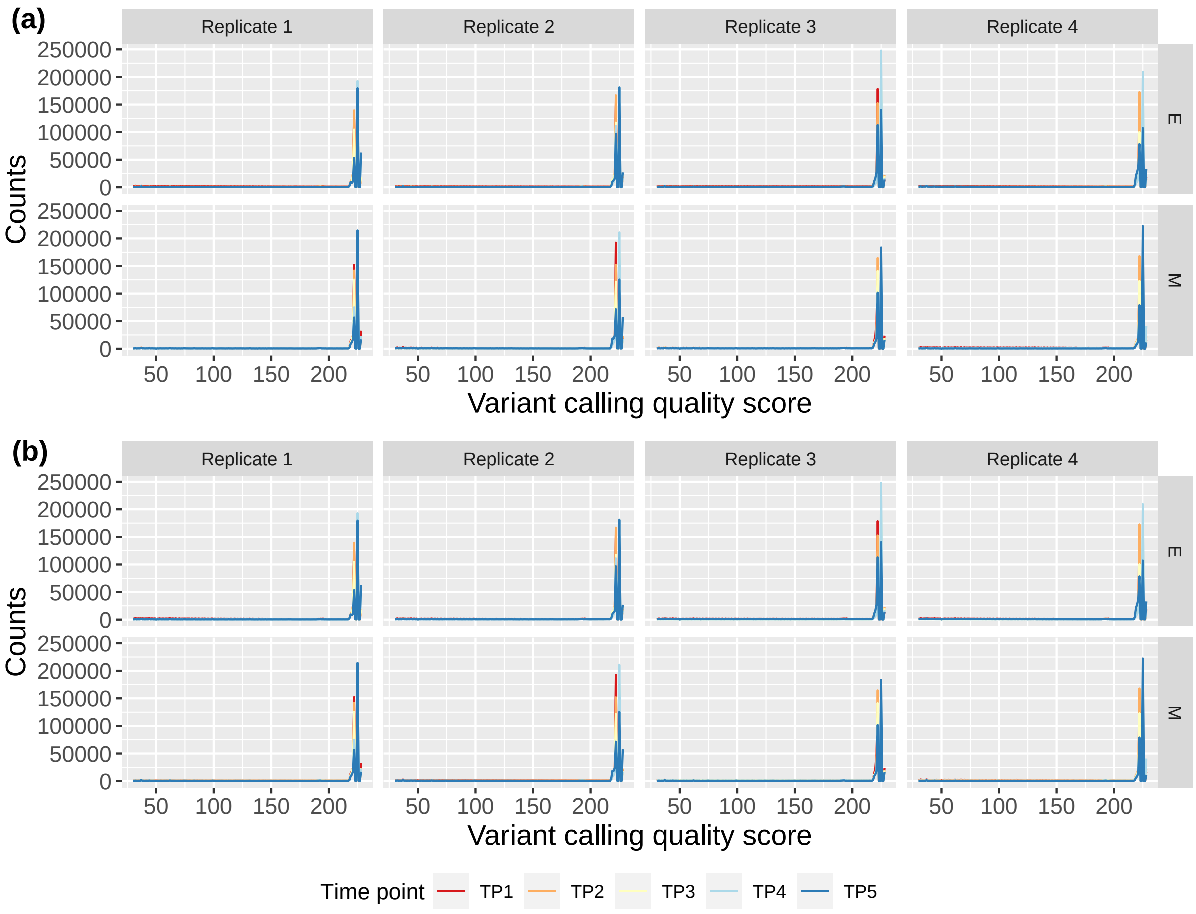


Figure S3: **Variant calling phred quality score per time point for each replicate.** (a) shows quality score distributions at the genome level and (b) at the X chromosome level assemblies. Each row corresponds to a different treatment - E on top row and M on bottom - and each column to an individual replicate. Time points are visible by differently coloured lines as per legend at the bottom of the graph.

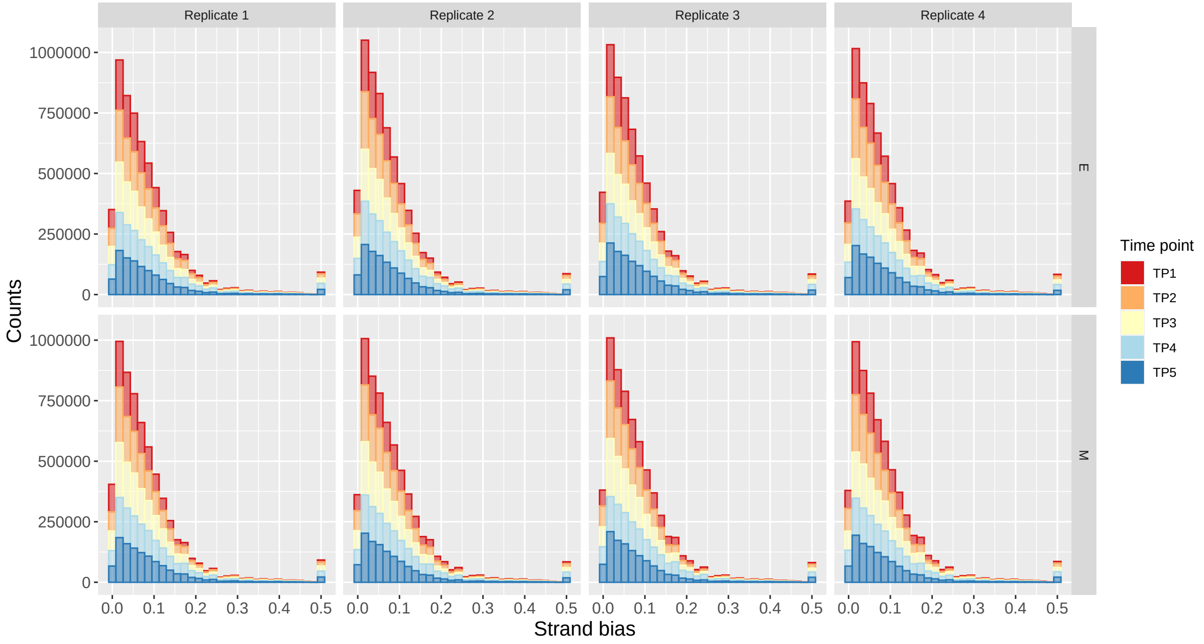


Figure S4: **Strand bias after filtering per time point for each of the four experimental replicates.** E populations are on the top tow, and M on the bottom. Each column corresponds to an individual replicate. Time point distributions are coloured differently as per side legend.

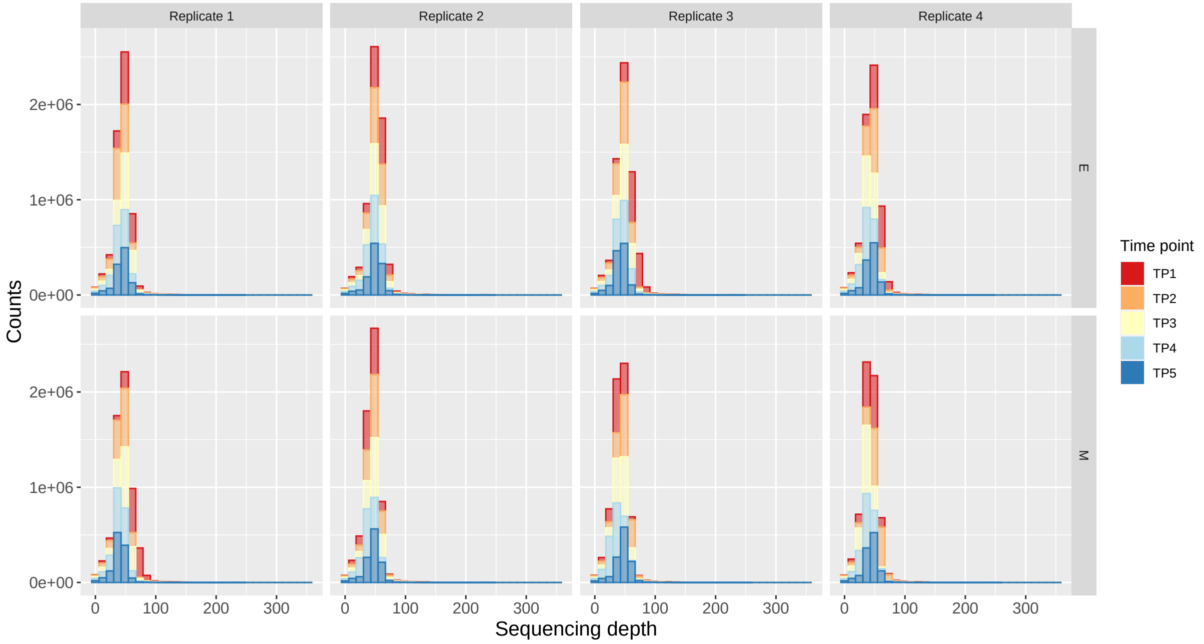


Figure S5: **Sequencing depth distribution per time point for all variants called and retained after filtering.** Replicates can be found in columns, and treatments in rows (E: top, M: bottom). Distributions for each time point were coloured differently as per side legend.

| Treatment | Time point | Unfiltered | Filtered round #1 (vs unfiltered) | | Both callers (vs filtered round #1) | | Filtered round #2 (vs both callers) | | Freebayes (vs unfiltered) | | Assembly level |
|---|---|---|---|---|---|---|---|---|---|---|---|
| M | TP1 | 2,787,319 | 2,477,272 | (-310,046.25) | 1,466,325 | (-1,010,947.75) | 1,166,114.75 | (-300,209.75) | 3,156,979 | (+369,660.75) | whole genome |
| M | TP2 | 2,491,572 | 2,303,865 | (-187,707) | 1,435,954 | (-867,911.25) | 1,387,945.25 | (-48,008.5) | 2,305,271 | (-186,300.75) | whole genome |
| M | TP3 | 2,351,323 | 2,182,542 | (-168,781.5) | 1,384,611 | (-797,930.75) | 1,358,823.50 | (-25,787.5) | 2,117,929 | (-233,394.75) | whole genome |
| M | TP4 | 1,967,133 | 1,847,275 | (-119,857.75) | 1,240,078 | (-607,197.75) | 1,204,845.00 | (-35,232.5) | 1,632,634 | (-334,499) | whole genome |
| M | TP5 | 1,948,784 | 1,842,966 | (-105,817.75) | 1,209,587 | (-633,379.25) | 1,189,898.50 | (-19,688.5) | 1,754,641 | (-194,142.75) | whole genome |
| E | TP1 | 2,843,322 | 2,547,546 | (-295,776) | 1,498,669 | (-1,048,876.25) | 1,214,163.25 | (-284,506) | 2,910,488 | (+67,166.5) | whole genome |
| E | TP2 | 2,453,723 | 2,279,650 | (-174,072.75) | 1,430,774 | (-848,876) | 1,386,282.50 | (-44,491.5) | 2,279,557 | (-174,165.75) | whole genome |
| E | TP3 | 2,135,627 | 2,013,440 | (-122,186.5) | 1,301,792 | (-711,648) | 1,283,460.00 | (-18,332.25) | 1,861,553 | (-274,074.25) | whole genome |
| E | TP4 | 1,910,089 | 1,802,535 | (-107,553.75) | 1,210,829 | (-591,705.75) | 1,176,927.25 | (-33,901.75) | 1,573,896 | (-336,193) | whole genome |
| E | TP5 | 2,062,411 | 1,929,839 | (-132,571.5) | 1,242,848 | (-686,991.5) | 1,214,033.25 | (-28,814.5) | 1,979,512 | (-82,898.75) | whole genome |
| M | TP1 | 1,047,522 | 942,455 | (-105,067) | 565,788 | (-376,666.5) | – | – | 1,182,418 | (-134896) | X |
| M | TP2 | 925,122 | 852,754 | (-72,368.5) | 528,325 | (-324,429) | – | – | 900,689 | (+24433.25) | X |
| M | TP3 | 871,490 | 804,422 | (-67,068) | 506,213 | (-298209) | – | – | 832,029 | (+39461) | X |
| M | TP4 | 731,140 | 676,547 | (-545,92.75) | 450,819 | (-225728.5) | – | – | 654,219 | (+76920.5) | X |
| M | TP5 | 739,132 | 687,412 | (-517,19.75) | 445,051 | (-242361.75) | – | – | 726,722 | (+12410.25) | X |
| E | TP1 | 1,055,999 | 949,923 | (-106,076) | 560,418 | (-389505) | – | – | 1,100,105 | (-44105.5) | X |
| E | TP2 | 915,083 | 846,588 | (-68,494.5) | 530,034 | (-316553.75) | – | – | 897,880 | (+17202.25) | X |
| E | TP3 | 777,659 | 724,817 | (-52,841.5) | 465,100 | (-259717.25) | – | – | 733,010 | (+44648.25) | X |
| E | TP4 | 690,257 | 642,877 | (-47,379.25) | 430,364 | (-212513.75) | – | – | 611,128 | (+79128.25) | X |
| E | TP5 | 770,680 | 711,135 | (-59,544.75) | 452,685 | (-258450) | – | – | 810,071 | (-39391.25) | X |

Table S3: **Number of SNPs at different stages of parsing for M and E lines.** 'Unfiltered' are the average total number of SNPs at each time point called by bcftools before any parsing. Subsequent filtering for variants called with a quality score of at least 30 resulted in the 'Filtered round #1' column, where numbers in brackets are the number of SNPs lost in this parsing step. Only SNPs that were called by both bcftools and Freebayes were retained after the first filtering step - 'Both callers'. Variants lost here are in brackets. 'Filtered round #2' included keeping biallelic sites only, as well as retaining solely those variants that were called both in the bwa mem and the novoalign alignment. In first time point samples, only polymorphic sites with a MAF $\geq 0.025$ were kept. SNPs called by Freebayes are included for comparison ('Freebayes'). Figures in brackets here are the difference to 'Unfiltered' polymorphisms.

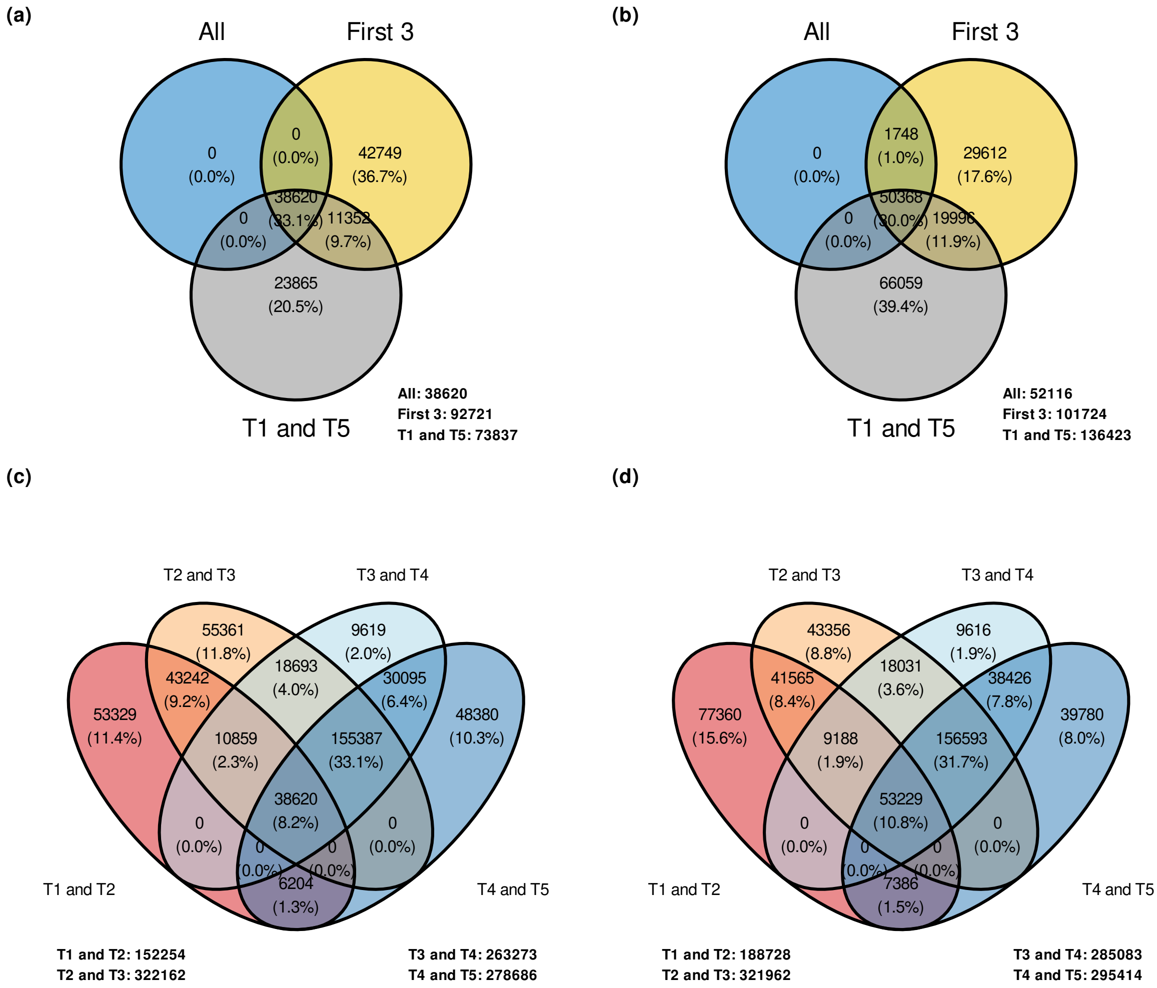


Figure S6: **Venn diagrams that compare the final number of SNPs analysed between different time point interval datasets.** Panels (a) and (b) show number of SNPs found in a five time point time series ('All'), a three time point time series ('First 3') or the first and last time points (T1 and T5) for M and E, respectively. Bottom panels (c) – M lines – and (d) – E lines – compare two time point intervals: T1 and T2, T2 and T3, T3 and T4, and T4 and T5.

| Interval | Treatment | Chromosome 2 | Chromosome 3 | Chromosome 4 | Chromosome X | Total |
|---|---|---|---|---|---|---|
| All time points | M | 11128 | 10537 | 6593 | 9807 | 38065 |
| First three | M | 25300 | 22547 | 18568 | 25191 | 91606 |
| T1T5 | M | 21084 | 17144 | 13186 | 21396 | 72810 |
| T1T2 | M | 43477 | 33217 | 30954 | 42911 | 150559 |
| T2T3 | M | 78318 | 58094 | 63784 | 119452 | 319648 |
| T3T4 | M | 68139 | 45920 | 51847 | 95260 | 261166 |
| T4T5 | M | 71413 | 45159 | 57190 | 102471 | 276233 |
| All time points | E | 12189 | 14021 | 11422 | 13707 | 51339 |
| First three | E | 25358 | 23347 | 23278 | 28438 | 100421 |
| T1T5 | E | 32883 | 26935 | 27344 | 47587 | 134749 |
| T1T2 | E | 48018 | 35596 | 38963 | 64042 | 186619 |
| T2T3 | E | 78890 | 54384 | 75011 | 111092 | 319377 |
| T3T4 | E | 70816 | 46865 | 68182 | 96712 | 282575 |
| T4T5 | E | 70666 | 47206 | 75381 | 99531 | 292784 |

Table S4: **Final number of SNPs per treatment for several time point intervals used for subsequent analyses.** 'All time points' and 'First three' correspond to the two time series we have analysed which have information on either 5 or 3 time points, respectively. T1T5, T1T2, T2T3, T3T4 and T4T5 are every interval combination of any two time points. Total SNP numbers for chromosomes 2, 3, 4 and X can be found in columns 3 to 6.

### 4 Supplement to diversity analysis

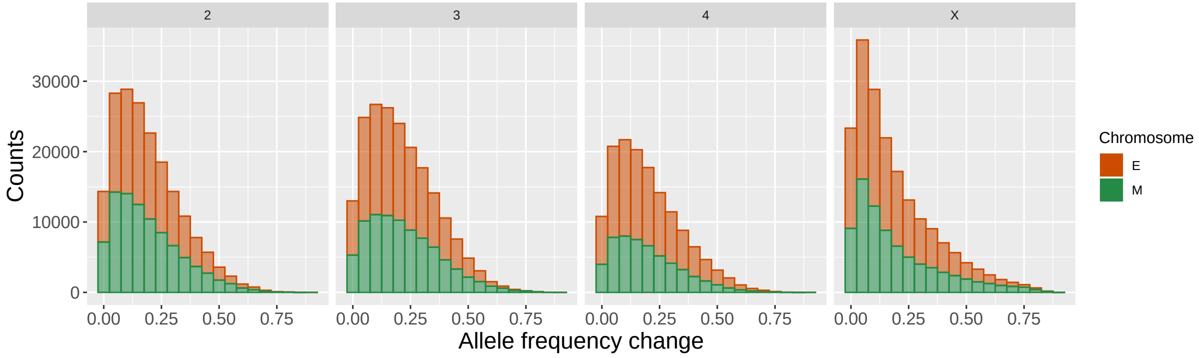


Figure S7: **Allele frequency change histograms in M (in green) and E (in orange) populations for each chromosome (columns).** These are calculated as the difference in allele frequency between first and last time point for each individual SNP.

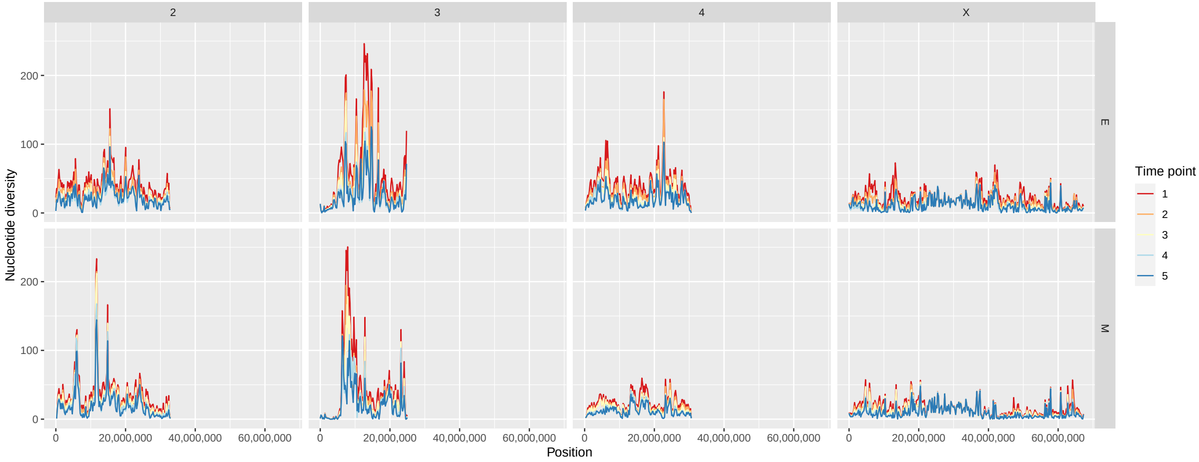


Figure S8: **Average nucleotide diversity, $\pi$, along the genome.** Columns correspond to chromosomes and rows to the two different treatments (top: E; bottom: M). Lines are coloured as to show variation across time. Averages were calculated across replicates for each 250k SNP window separately.

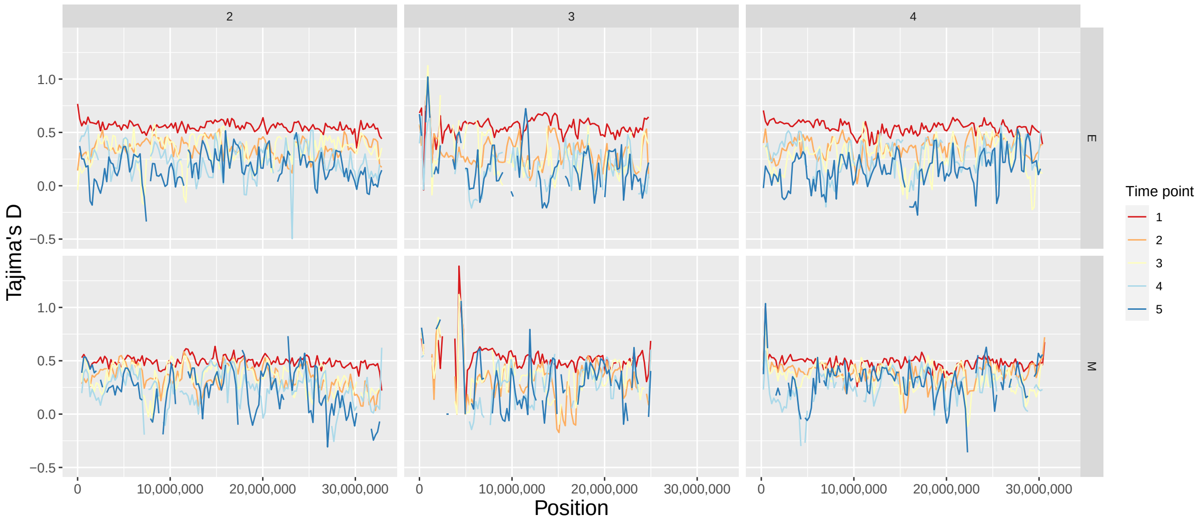


Figure S9: **Tajima's D estimates along chromosomes 2, 3 and 4 for E and M lines.** Rows correspond to the two different treatments and columns to chromosomes. Estimates were calculated in 250k SNP windows with Grenedalf (Czech and Exposito-Alonso, 2021). Lines are coloured per time point according to the side legend.

#### 5 Supplement to $N_e$ analysis

| Time interval | Median - M | Median - E |
|---|---|---|
| Overall | 151.0 (n = 100) | 159.2 (n = 223) |
| T1T2 | 90.0 (n = 212) | 85.7 (n = 308) |
| T2T3 | 68.2 (n = 246) | 111.9 (n = 239) |
| T3T4 | 73.8 (n = 131) | 102.9 (n = 112) |
| T4T5 | 134.8 (n = 107) | 145.8 (n = 116) |

Table S5: **Median genome-wide $N_e$ estimates for M and E lines at different time point intervals using intergenic SNPs only.** Medians were calculated using 1k intergenic SNP window estimates from all of the four experimental replicates. 'Overall' corresponds to $N_e$ estimates based on allele frequency changes between the first and last time point. The total number of windows considered in each replicate is found in brackets.

### Supplement to genome scan

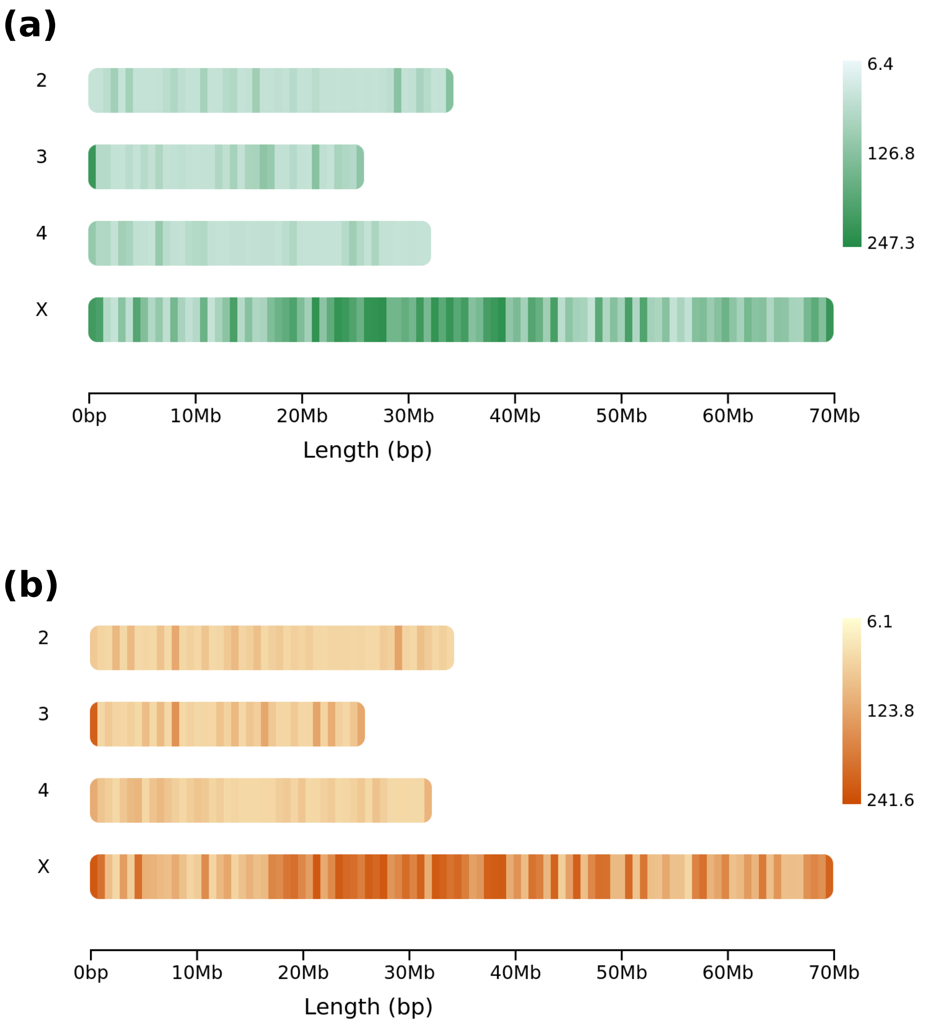


Figure S10: **Maximum coverage chromosome plots for (a) M and (b) E lines.** Each diagram represents the maximum coverage of any given interval in each of the four chromosomes analysed. M lines are coloured in green and E in orange. Highest coverage regions can indicate repetitive elements and are common in telomeres and centromeres. Data exclude chromosome 5.

| Treatment | Chromosome | Location | NCBI ID | FB ID | *D. mel* ortholog | Gene name | #top SNPs | Distance |
|---|---|---|---|---|---|---|---|---|
| E | 2 | Gene | 6896726 | FBgn0262041 | hdc | headcase protein | 1 | NA |
| E | 3 | Gene | 4805669 | FBgn0073138 | SmydA-5 | SET domain-containing protein SmydA-8 | 1 | NA |
| E | 3 | Gene | 4805666 | FBgn0080959 | Cyp4aa1 | cytochrome P450 4aa1 | 1 | NA |
| E | 3 | Gene | 6899100 | FBgn0246519 | Pgant9 | polypeptide N-acetylgalactosaminyltransferase 9 | 1 | NA |
| E | 3 | Gene | 6899112 | FBgn0246524 | Verprolin 1 | WAS/WASL-interacting protein family member 3 | 1 | NA |
| E | 3 | Gene | 6899088 | FBgn0263807 | Strn-Mlck | myosin light chain kinase | 2 | NA |
| E | X | Gene | 4815279 | FBgn0075585 | fzr | fizzy-related protein homolog | 3 | NA |
| E | X | Gene | 4814581 | FBgn0078871 | forked | espin | 3 | NA |
| E | X | Gene | 4815373 | FBgn0080302 | CG7378 | dual specificity protein phosphatase 3 | 1 | NA |
| E | X | Gene | 4814961 | FBgn0081931 | CG9657 | sodium-coupled monocarboxylate transporter 1 | 1 | NA |
| E | X | Gene | 4814480 | FBgn0243547 | Atg5 | autophagy protein 5 | 4 | NA |
| E | X | Gene | 6901434 | FBgn0247979 | dpr8 | zwei Ig domain protein zig-8 | 5 | NA |
| M | X | Gene | 6902114 | FBgn0244416 | CG32532 | homeobox protein Hmx | 1 | NA |
| E | 3 | Intergenic region | 4805672 | FBgn0073140 | CCHa2-R | neuropeptide CCHamide-2 receptor | 1 | -421 |
| E | 3 | Intergenic region | 6899092 | FBgn0245480 | resilin | pro-resilin | 2 | -579 |
| E | 4 | Intergenic region | 4816335 | FBgn0077513 | CG3528 | cilia- and flagella-associated protein 299 | 1 | -9236 |
| E | X | Intergenic region | 4815048 | FBgn0074111 | unc-119 | protein unc-119 homolog | 1 | 2760 |
| E | X | Intergenic region | 4814454 | FBgn0080287 | Flacc | fl(2)d-associated complex component | 1 | 1833 |
| E | X | Intergenic region | 4814479 | FBgn0081927 | brinker | J domain-containing protein DDB_G0295729 | 1 | -3996 |
| E | X | Intergenic region | 6901756 | FBgn0249747 | CG15034 | uncharacterised protein | 1 | -516 |
| E | X | Intergenic region | 6901755 | FBgn0249791 | - | antigen 5 like allergen Cul n 1 | 1 | 1613 |

Table S6: **Common genes amongst top scoring variants in this study and Wiberg et al. (2021).** This includes data on genes where significant variants are located in or genes near top SNPs. Each gene is described by an NCBI ID, a FlyBase ID, a *D. melanogaster* ortholog, the gene's name, how many top SNPs mapped on to the gene or the distance between any intergenic SNPs and the nearest gene.

| Chromosome | Treatment | # top SNPs | Gene NCBI ID | Gene FB ID | *D. mel* ortholog | Gene name |
|---|---|---|---|---|---|---|
| X | E | 7 | 4813031 | FBgn0076699 | Lmx1a | LIM homeobox transcription factor 1-beta |
| X | E | 9 | 4813494 | FBgn0076929 | SpoCk | calcium-transporting ATPase type 2C member 1 |
| X | E | 11 | 4813557 | FBgn0076932 | neuromusculin | hemicentin-1 |
| X | E | 6 | 4814416 | FBgn0077610 | Fas2 | fasciclin-2 |
| X | E | 5 | 4814942 | FBgn0079214 | Nep1 | neprilysin-1 |
| 3 | E | 5 | 6899052 | FBgn0263814 | jing | zinc finger protein jing |
| 3 | E | 5 | 6899426 | FBgn0250039 | luna | Krueppel-like factor luna |
| X | E | 23 | 6901139 | FBgn0243670 | CG43867 | uncharacterised protein |
| X | E | 5 | 6901434 | FBgn0247979 | dpr8 | zwei Ig domain protein zig-8 |
| X | E | 7 | 6901448 | FBgn0247929 | Sh | potassium voltage-gated channel protein Shaker |
| X | E | 7 | 6901459 | FBgn0247897 | CG5921 | uncharacterised protein |
| X | E | 5 | 6901716 | FBgn0245264 | - | voltage-dependent T-type calcium channel subunit alpha-1G |
| X | E | 9 | 6901746 | FBgn0245096 | SK | small conductance calcium-activated potassium channel protein |

Table S7: **Genes with the most significant variants.** Each gene is described by the number of top SNPs located in said gene, an NCBI ID, a FlyBase ID, a *D. melanogaster* ortholog and the gene's name.
